# A unified circuit model of attention: Neural and behavioral effects

**DOI:** 10.1101/2019.12.13.875534

**Authors:** Grace W. Lindsay, Daniel B. Rubin, Kenneth D. Miller

## Abstract

Selective visual attention modulates neural activity in the visual system in complex ways and leads to enhanced performance on difficult visual tasks. Here, we show that a simple circuit model, the stabilized supralinear network, gives a unified account of a wide variety of effects of attention on neural responses. We replicate results from studies of both feature and spatial attention, addressing findings in a variety of experimental paradigms on changes both in firing rates and in correlated neural variability. Finally, we expand this circuit model into an architecture that can perform visual tasks—a convolutional neural network—in order to show that these neural effects can enhance detection performance. This work provides the first unified mechanistic account of the effects of attention on neural and behavioral responses.

## 1. Introduction

When an animal knows in advance what features or locations in the visual scene will be relevant for completing its goals, selective top-down attention can be deployed. This attention has been shown to have a powerful modulatory effect on both task performance and neuronal responses, and changes in the latter can often be powerful predictors of the former (Ress et al., 2000).

Numerous specific impacts of attention on neural activity have been identified, including changes in firing rates, trial-to-trial variability, and noise correlations (Cohen and Maunsell, 2009; Treue and Martinez Trujillo, 1999; Treue and Maunsell, 1999). Looking at the impact of attention on tuning curves, attention to a preferred stimulus is known to scale up the responses to all stimuli; conversely, attention to a non-preferred stimulus scales responses down (Martinez-Trujillo and Treue, 2004). This enhancement has been shown to be a largely multiplicative increase in neuronal gain (Treue and Martinez Trujillo, 1999). A similar percentage change occurs in the firing rates of excitatory and inhibitory neurons (Mitchell et al., 2007).

Many of attention’s impacts on firing rates can be understood in the context of the normalization model of attention (Boynton, 2009; Ghose, 2009; Lee and Maunsell, 2009; Reynolds and Heeger, 2009). This model builds off the canonical computation of normalization observed in multiple places in the visual system as well as other brain areas (Carandini and Heeger, 2012). In the absence of attention, a neuron’s firing rate can be predicted by a divisive normalization equation: stimuli with the preferred features and in the classical receptive field of the neuron form the numerator (known as the “stimulus drive”), and the denominator is a function of a less-selective suppressive drive that includes surround locations and non-preferred features. Under the normalization model of attention, attention provides a biasing effect that amplifies the drive coming from the attended stimulus.

This model captures how attention can, when two stimuli are present, shift responses to those of the attended stimulus alone. For example, when a preferred and non-preferred stimulus are both presented to the receptive field of a V4 neuron, the cell’s response is intermediate between the responses evoked by each stimulus alone. By attending to either the preferred or non-preferred stimulus, the response is shifted towards the response evoked by the attended stimulus alone (Reynolds and Desimone, 2003). Similarly, attention to a stimulus in the suppressive surround of a V4 neuron increases the suppression induced, whereas attention to the center reduces the suppression (Sundberg et al., 2009). The normalization model of attention also captures how attention increases contrast gain or response gain, respectively, depending on whether the attention is over a larger or smaller cortical area than the stimulus input (Reynolds and Heeger, 2009).

Beyond changes in firing rates described by the normalization model of attention, attention also decreases trial-to-trial variability and noise correlations across neuron pairs (Cohen and Maunsell, 2009; Mitchell et al., 2007).

The normalization model is a phenomenological description: a mathematical description of responses with no underlying mechanistic model. While circuit models of some attentional phenomena have been presented (Compte and Wang, 2006; Miconi and VanRullen, 2016; Sajedin et al., 2019), there exists no unified circuit model that captures both a broad range of neural effects and the behavioral effects of attention. Such a model would provide a mechanistic test bed to probe anticipated effects of experimental manipulations and so guide future experiments studying attention. Here we present such a unified circuit model of attention.

We have previously shown that a simple model of cortical circuitry—known as the stabilized supralinear network (SSN) (Ahmadian et al., 2013)—can account for a wide set of phenomena described by normalization, including feature normalization and surround suppression and their nonlinear dependencies on contrast (Rubin et al., 2015). It also accounts for the suppression of correlated variability by a stimulus (Hennequin et al., 2018). The network assumes expansive or supralinear input/output functions for the individual units. As described in (Ahmadian and Miller, 2019; Ahmadian et al., 2013; Rubin et al., 2015), this causes effective synaptic strengths between units—which are proportional to the postsynaptic neuron’s gain (its change in firing rate for a given change in input) – to grow with increasing postsynaptic activation. The growth of excitatory-to-excitatory effective connections leads to potential instability, but with sufficiently strong feedback inhibition the network remains stable. However, this stabilization occurs through the network dynamically “loosely balancing” its inputs, so that the recurrent input largely cancels the feedforward input, leaving a residual net input that grows sublinearly as a function of the feedforward input. (The balancing is “loose” because the residual input after cancellation is comparable in size to the factors that cancel, Ahmadian and Miller, 2019.) This cancellation of feedforward input through increasingly strong inhibitory stabilization leads to the normalization and suppression of variability just described.

As a result of its strong recurrent excitation stabilized by strong feed-back inhibition, the SSN exhibits “balanced amplification” (Hennequin et al., 2018; Murphy and Miller, 2009): small inputs biased toward either excitatory (or inhibitory) cells drive large increases (or decreases) in both excitatory and inhibitory firing rates. We hypothesized that attentional modulation acts through the same balanced amplification and recurrent “loose balancing” mechanisms that implement feature normalization and surround suppression. Here we show that this model can indeed account for a strikingly large range of the neural effects of attention observed in visual cortex.

Can the attentional neural modulations of the SSN yield attentional enhancement of performance? To address this, we build on previous work (Lindsay and Miller, 2018). In that work, we used a deep convolutional neural network (CNN) as a model of the visual system to show how neural changes associated with attention enhance performance on a challenging visual detection task. In that model, attention was implemented by changing the gain of units “by hand” according to their relationship to the attended stimulus. Here, we instead put our circuit model of attentional modulation into the CNN architecture, demonstrating that a biological circuit implementation of attentional neural modulation can induce attentional modulations of behavior like those observed biologically. This model (dubbed the SSN-CNN) replicates both the neural impacts of attention and the performance enhancements.

## 2. Results

We employ four instantiations of the SSN model to replicate the neural effects of attention. The details of all of these models have been described previously (Rubin et al., 2015), and are included in the Methods section. All four models feature strongly recurrently connected excitatory and inhibitory neurons with a supralinear neuronal input-output nonlinearity. The four models differ only in the dimension of stimulus space over which the neurons are arranged and the spatial arrangement and strengths of the connections between neurons. In the simplest model, we consider a single pair of excitatory (E) and inhibitory (I) neurons (Figure 1A). In the first of the larger models, pairs of E and I neurons are arranged around a ring, with position on the ring corresponding to a periodic preferred feature such as stimulus orientation or direction, with all cells having a similar retinotopic receptive field (RF) position (Methods 4.1.1, Figure 1B). Alternatively, E-I pairs are arranged on a line, with position on the line interpreted as retino-topic RF position of the cells, which otherwise have similar preferred stimulus features (Methods 4.1.2, Figure 8C). The final, most complex model has a 2-dimensional representation of retinotopic space on which is superimposed a map of preferred features, taken to have the structure of a map of preferred orientations in V1 (Kaschube et al., 2010). In this model, neurons make connections as probabilistic functions of difference in stimulus preference over the three dimensions of stimulus quality: two spatial dimensions and the stimulus feature dimension (Methods 4.1.3). Because connectivity is probabilistic, this model has cell-to-cell diversity in connections, allowing us to study cell-to-cell diversity in responses.

**Figure 1:**
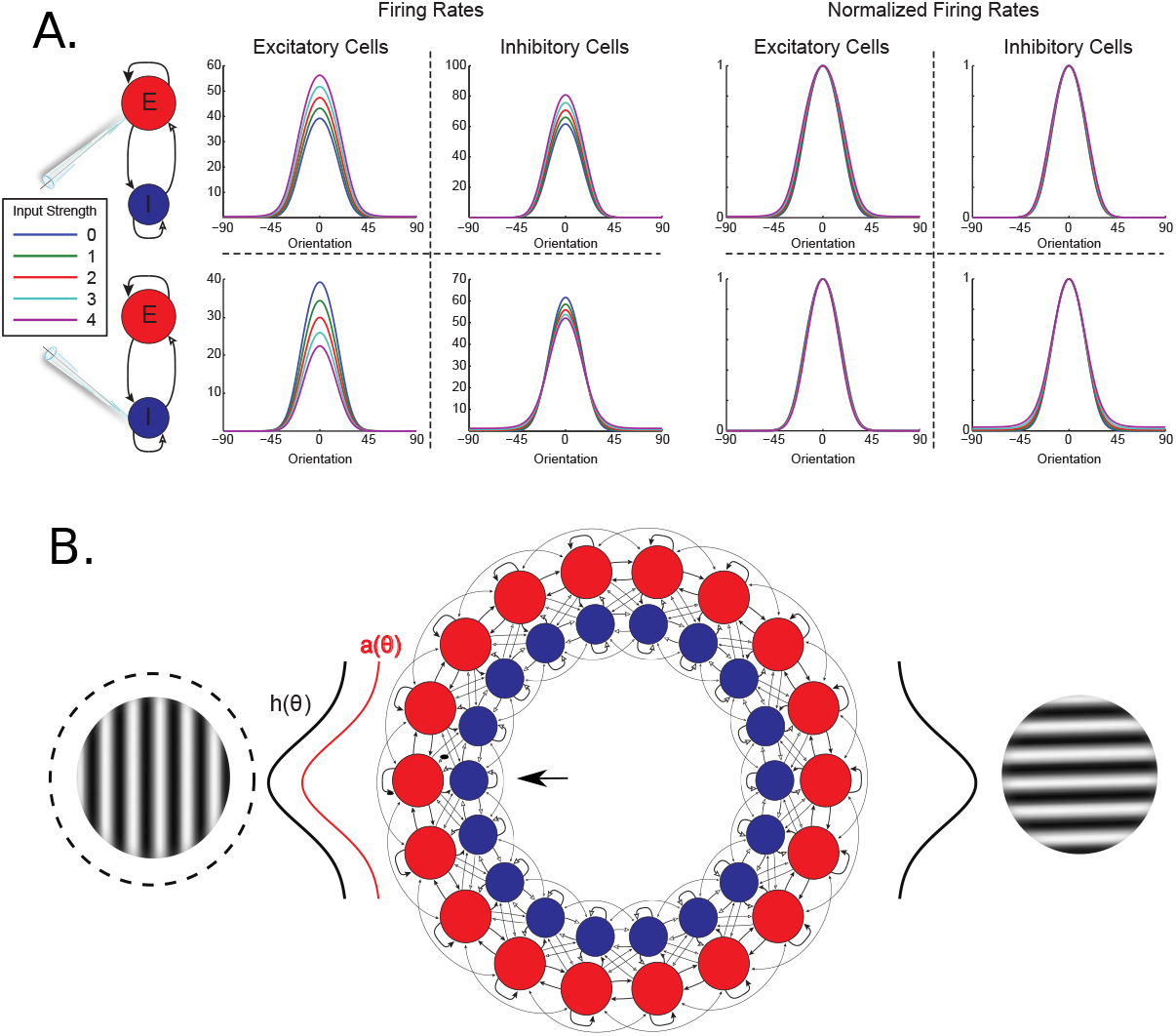
A.) Expansive nonlinearity and balanced amplification yield multiplicative scaling. We consider a simple two-unit nonlinear SSN model, with one excitatory (E) cell and one inhibitory (I) cell (Methods 4.1.4). We drove both cells with a series of feedforward inputs, whose strengths varied as a function of “orientation” to generate “tuning curves”. While driving the cells with this feedforward input, an additional constant input of one of four varying strengths (indicated by color legend at left) was added to either the E or the I cell. With increasing input to the E cell, both E and I rates are scaled up, whereas with increasing input to the I cells, both E and I rates are scaled down. Normalizing each curve by its maximum reveals that the gain change is almost exclusively multiplicative. B.) A ring model of attention. The ring model represents different features (*e.g*., preferred orientation) at a single location in visual space. At each location on the ring, a pair of excitatory (red) and inhibitory (blue) cells exist. Oriented stimuli are modeled as Gaussians centered at a particular location on the ring (black curves). Attention to one of the stimuli (indicated by dashed circle around it) is modeled as an additional Gaussian input biased towards the excitatory subpopulation at the center of the locus of attention (red curve). In this example, recording from the E-I pair indicated with the arrow would correspond to the cyan line in Figure 2A

In our models, the inputs to the model cortex are assumed to sum linearly, so that all nonlinear behavior arises from cortical processing. However, we note that the suppression of response to a preferred orientation by simultaneous presentation of an orthogonal orientation or “mask” (“cross-orientation suppression”) in V1 is largely mediated by nonlinear changes in the pattern of thalamic firing induced by the mask, rather than by nonlinear V1 integration (Li et al., 2006; Priebe and Ferster, 2006) (although there is a component mediated by V1 as shown by suppression arising when the two stimuli are presented to different eyes, Sengpiel and Vorobyov, 2005). We typically refer to different competing stimuli presented within an RF as “orientations”, but this should be understood to model cortical processing given linear summation of inputs induced by two stimuli, rather than the literal phenomenon of V1 cross-orientation suppression.

In all instantiations, attention is modeled as a small additional excitatory input given to the excitatory cells within the specified locus of attention. As a secondary test, we also re-ran all simulations with attention instead modeled as a small inhibitory input to the inhibitory cells (resulting in a disinhibition of locally-connected excitatory cells). Results were qualitatively similar, with a few notable exceptions discussed below.

To investigate the impact of neural activity changes on performance, we also incorporated one of these circuit models—the ring model—into a convolutional neural network architecture (Methods 4.3). This allowed us to demonstrate that the application of attention to our circuit model can increase performance on a challenging visual detection task.

### 2.1. Basic mechanism of the model

The balanced amplification model (Murphy and Miller, 2009) demonstrates that in a network with strong recurrent connectivity, small changes in the difference between E and I activity can drive large changes in the sum of the activity. Previously, we have used this mechanism to produce models of contextual modulation that capture the experimental observation that, during surround suppression, both E and I firing rates are suppressed (Ozeki et al., 2009). Within a locus of attention, however, the opposite effect is observed: both E and I firing rates are enhanced (Mitchell et al., 2007).

In a network wherein neurons are described by a supralinear nonlinearity, a bias in the input towards E or I shifts the responses of both cells up or down (respectively), and the resulting change can be almost exclusively multiplicative (Figure 1A). Thus we hypothesize that this simple, intrinsic form of amplification may be sufficient to account for the observed effects of attention on visual cortical circuits. We now incorporate this simple E-I pair into a broader recurrent circuit and consider several recent experimental results on attention in visual cortex.

### 2.2. Attention influences stimulus interactions

#### 2.2.1. Impact of feature attention

In several regions of visual cortex, attention to one of multiple stimuli presented within the receptive field of a neuron can shift the response of that neuron towards the response evoked by the attended stimulus alone. This was shown by Reynolds and Desimone (2003), who probed the responses of V4 neurons with preferred and non-preferred stimuli, presented either alone or together in the receptive field of a single neuron. They found that in the simultaneous presentation condition, attending to a non-preferred stimulus caused a relative suppression compared to an attend-away condition, whereas attending to the preferred stimulus boosted the response. To simulate this experiment, we recorded the response of a cell to a strong stimulus of preferred orientation in the ring model (for details of attention experiments see Methods 4.2). We then added a non-preferred stimulus at the orthogonal orientation to the ring (schematized in Figure 1B) and systematically varied the strength of this “probe” stimulus. As expected, the addition of the non-preferred probe was always suppressive, and with increasing probe strength suppression was increased (Figure 2A, blue line). We then repeated the same test with attention directed either towards the preferred stimulus (cyan) or the probe stimulus (green). When attention was directed towards the preferred stimulus, the amount of suppression was decreased. When attention was directed to the probe stimulus, suppression was enhanced.

**Figure 2:**
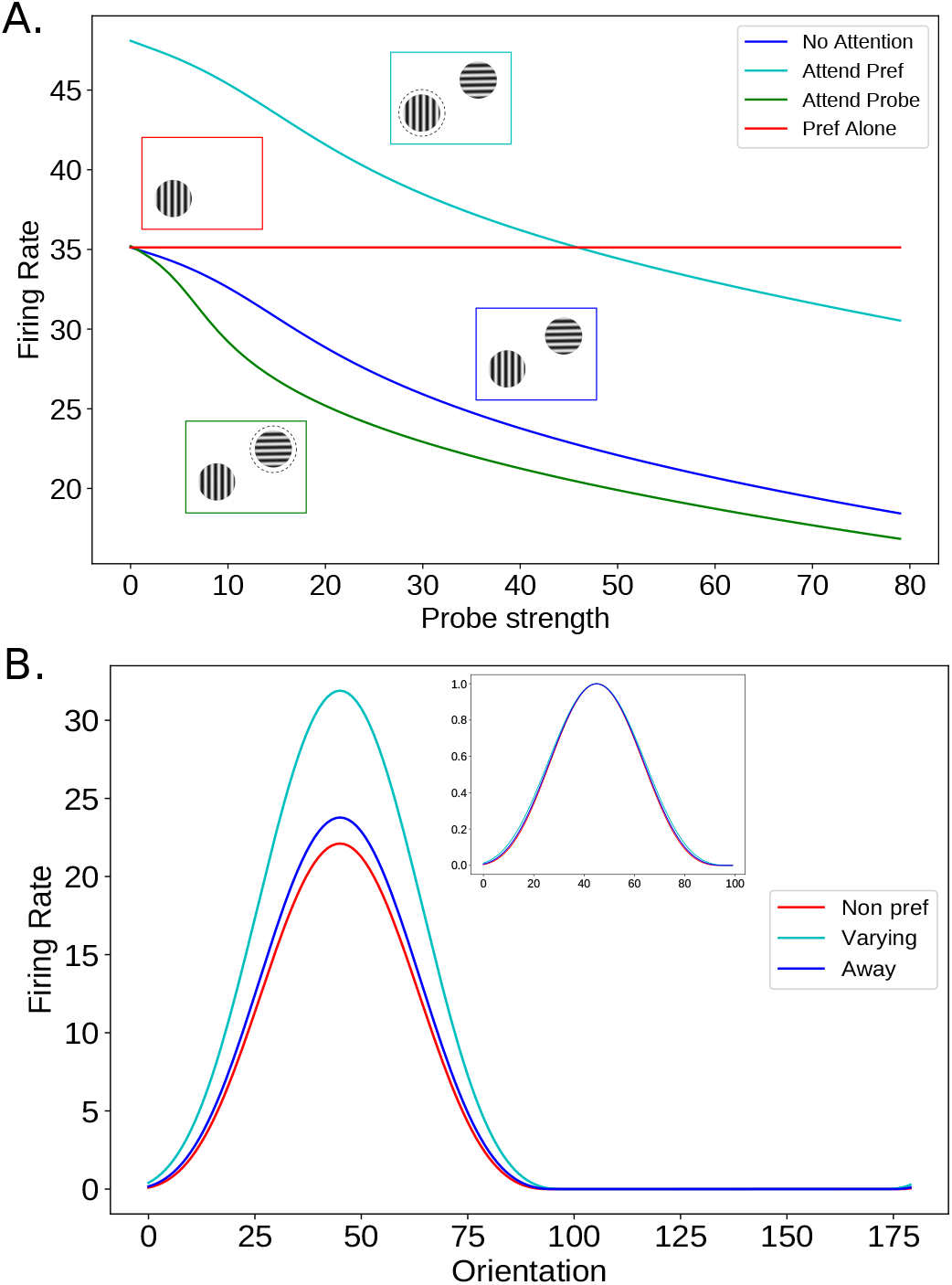
A.) Attention enhances the suppressive effect of non-preferred stimuli. A stimulus of preferred orientation was shown to a cell in the ring model. An orthogonally oriented stimulus was presented along with the preferred stimulus, and the strength of the non-preferred “probe” was varied (blue line). The test was then repeated with attention (indicated by dashed circle around stimulus) directed towards either the preferred stimulus (cyan) or the probe stimulus (green). When attention was directed towards the preferred stimulus, suppression was decreased. When attention was directed to the probe stimulus, suppression was enhanced. B.) Attention scales tuning multiplicatively. In the presence of a non-preferred probe stimulus, we varied the orientation of a test stimulus between 0° and 180°, while recording from the cell at 45° and attending either to the non-preferred probe (red), the varying stimulus (cyan), or away (blue). Attention produced an almost exclusively multiplicative change in response. Normalized responses are shown in the inset. There was virtually no change in tuning width, as observed experimentally (Treue and Martinez Trujillo, 1999).

In a related experiment, Treue and Martinez-Trujillo (1999) recorded from a neuron in area MT while presenting two stimuli to the neuron’s receptive field. One of the stimuli was always moving in a non-preferred direction, while the direction of the other stimulus was systematically varied. Compared to an attend-away condition, responses of MT neurons were relatively suppressed at all stimulus directions when attention was directed towards the non-preferred stimulus, but relatively enhanced when attending towards the varying stimulus. We find the same result if we repeat this test in our ring model (Figure 2B). Like Treue and Martinez-Trujillo (1999), the change we observe occurred without a substantial change in the width of tuning, indicating a mainly multiplicative scaling (Figure 2B, inset).

Note that in Figures 2A and 2B the same strength of attention is applied in all circumstances, however attention applied to a non-preferred stimulus has a weaker impact on firing rates. In our model, attention applied to a cell’s preferred stimulus results in additional excitatory input directly to the cell in question. Attention to an orthogonal stimulus only impacts the recorded cell indirectly through recurrent connections, leading to a weaker effect. Experimentally, the magnitude of firing rate changes has been found to be weaker when attention is applied to a non-preferred stimulus compared to a preferred one (Treue and Maunsell, 1999).

#### 2.2.2. Correlation between feature attention and normalization

Several groups have considered the mechanistic relationship between attention and cortical normalization (Bloem and Ling, 2019; Lee and Maunsell, 2009; Ni et al., 2012; Reynolds and Heeger, 2009). In a study exploring the variability in the strength of attentional modulation, Ni and collegues demonstrated that neurons vary in the degree to which their responses are normalized by the presence of an orthogonal, non-preferred stimulus in the receptive field. They further show that the degree of normalization a cell demonstrates (or in their terminology, the broadness of the “tuning” of normalization – quantified by a normalization modulation index) is highly correlated with the extent to which attention modulates the response to the cell. To simulate this experiment, we employed our 2-D model of visual cortex designed to reproduce both the mean effects as well as a realistic degree of variability in responses. In this simulation, excitatory cells were selected at random from the population. For each cell, a high contrast stimulus of preferred orientation was presented. An orthogonal stimulus of the same size, position, and strength (the “null” stimulus) was then presented, and then the preferred and orthogonal stimuli were presented together. The firing rate response in each of the three stimulus conditions was recorded, and the Normalization Modulation Index was calculated (see Methods 4.2. An NMI of 0.33 corresponds to averaging of the two stimuli, whereas an NMI of 0 is considered a “winner take all” response (the response to the pair is the same as the response to the preferred stimulus alone). In the terminology of Ni et al., cells with highly tuned normalization have an NMI closer to 0 (Ni et al., 2012). The paired presentations were then repeated (showing both preferred + null together) with attention directed towards either the preferred or null stimulus. Attention was applied to the E cells in the position, size, and orientation of either the preferred or null stimulus. An Attentional Modulation Index was then calculated. As was observed experimentally, there is a wide range of NMIs and AMIs, and the NMI and AMI of cells are highly correlated (Figure 3). A similar correlation was observed when inhibitory rather than excitatory cells were studied (not shown).

**Figure 3:**
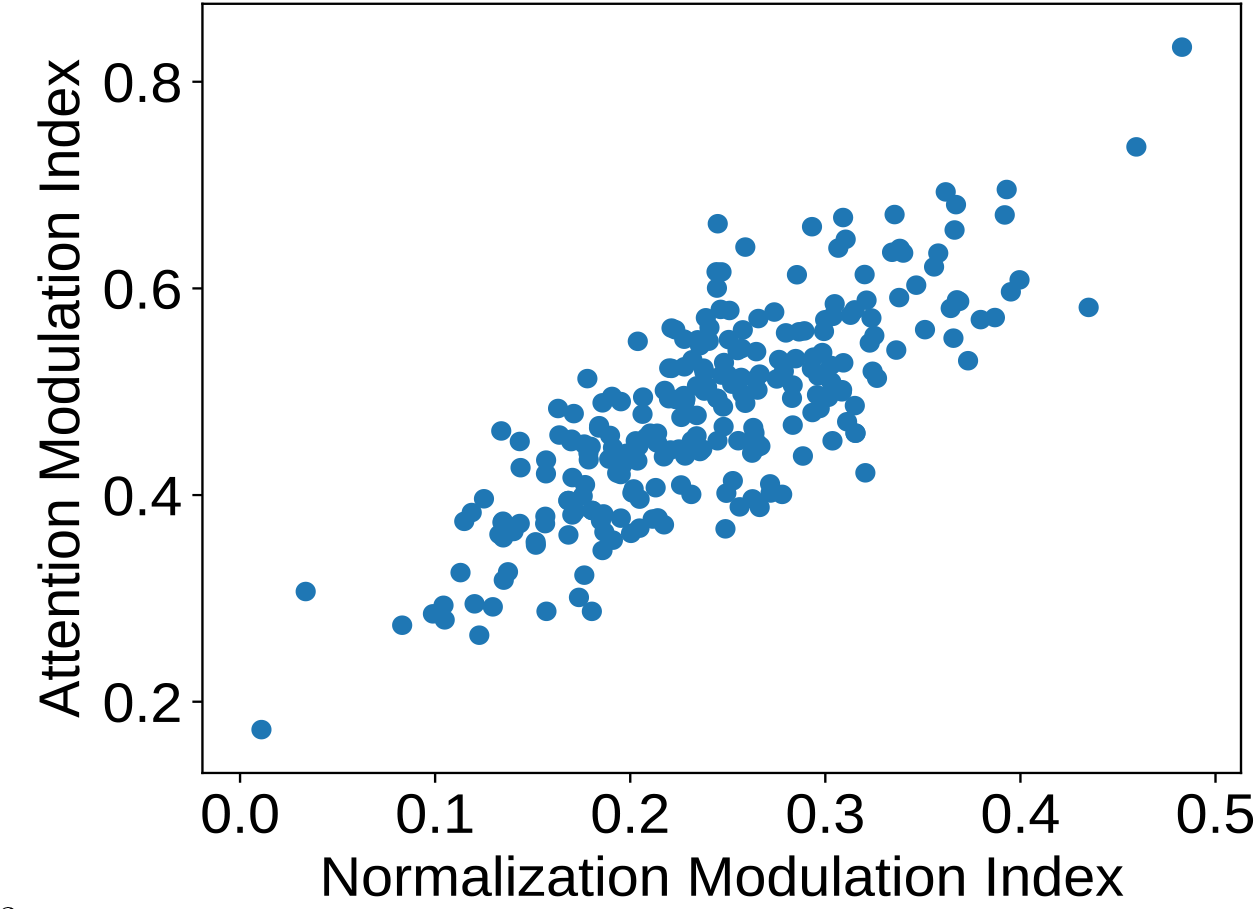
Normalization strength and attentional modulation are positively correlated. Normalization Modulation Indices are plotted against the Attention Modulation Indices for all 250 cells sampled from the 2-D model. Correlation coefficient: 0.80. See text for details.

This correlation was not induced simply by variability in the strength of excitatory or inhibitory connectivity received by cells, because in this model the excitatory input and inhibitory input to each cell are separately scaled so that each cell of a given type (excitatory or inhibitory) receives identical summed strength of excitatory input and of inhibitory input 4.1.3. Furthermore, the NMI or AMI were not correlated to the number of excitatory or number of inhibitory inputs received by a cell (not shown).

#### 2.2.3. Impact of spatial attention

The previously discussed experiments studied the response of neurons to pairs of stimuli presented within the same receptive field. However, attention has also been shown to modulate the effect of stimuli presented in the receptive field surround. Sundberg et al. (2009) found that in V4, the strength of surround suppression could be either increased or decreased by attending specifically to the surround or center stimulus. To simulate this experiment, we next employed our line model used to simulate spatial contextual interactions. Pairs of E and I cells are arranged along a one-dimensional lattice representing an axis of retinotopic space, with recurrent excitatory connections that decrease as a function of retinotopic/cortical distance. A stimulus was presented to the cell in the center of the lattice, in the presence of a suppressive surround stimulus. Attention was then directed to either the center or surround stimulus. Attention to the center decreased the strength of surround suppression (pushing firing rates towards those when the stimulus is presented alone), while attention to the surround enhanced surround suppression (Figure 4A).

**Figure 4:**
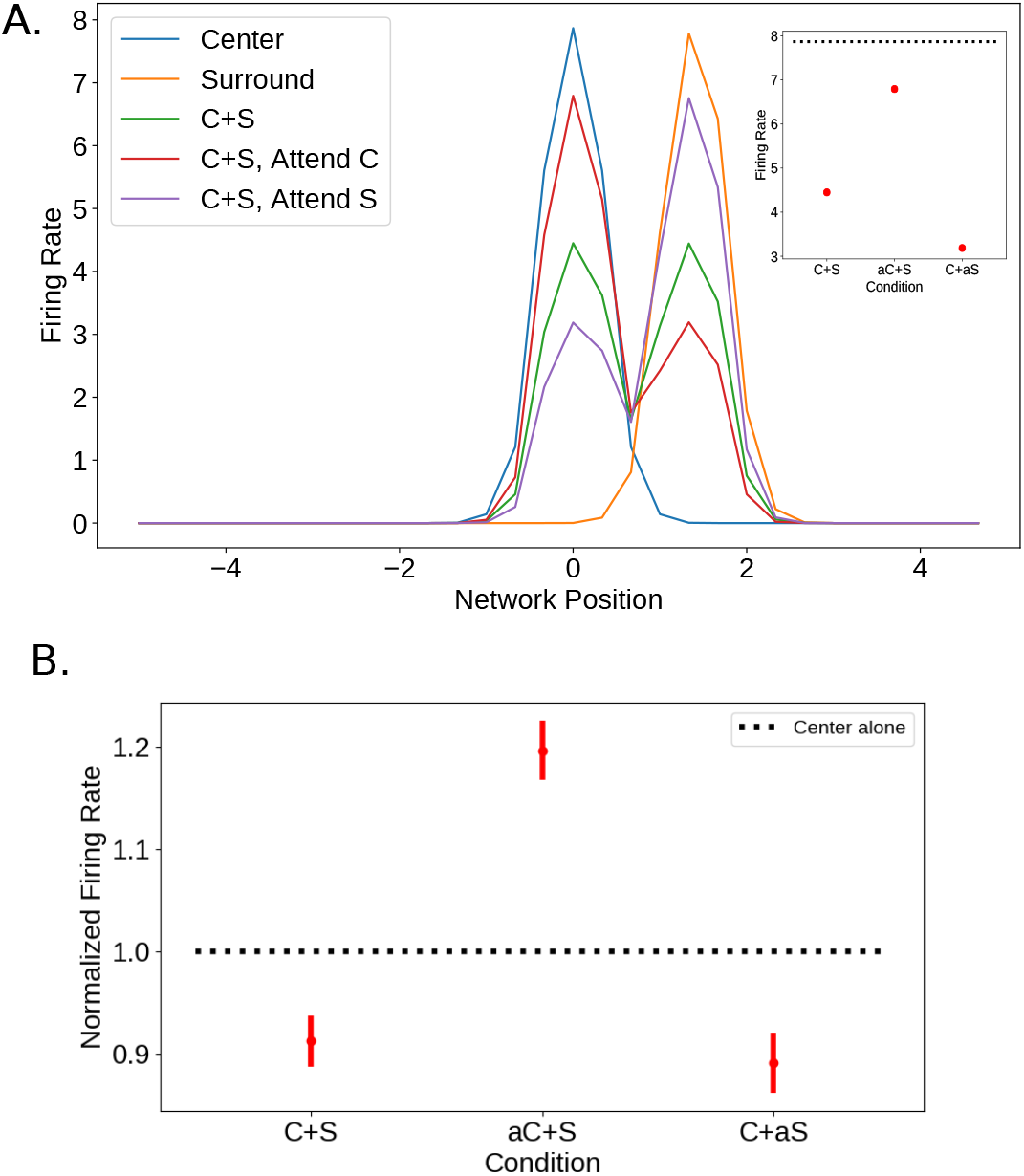
A.) Attention modulates the strength of surround suppression. A stimulus was shown in the receptive field of the neuron at position 0. A stimulus of equal strength and size was then placed in the surround, and the response was recorded from neurons in the vicinity. Attention was then directed either to the center or surround stimulus. In the main figure, the E cell activity across the network is shown in response to the center stimulus alone, the surround stimulus alone, the center and surround stimuli shown together, the center and surround stimuli with attention directed towards the center, and the center and surround stimuli with attention directed towards the surround. The inset demonstrates the activity at the center E cell – the dashed line is the response to the center stimulus alone, and the three dots show the response to the center and surround presented together, either with no attention, with attention directed towards the center, or with attention directed towards the surround. B.) Attention modulates the strength of surround suppression in the large scale model. A stimulus of preferred orientation was shown to a randomly selected cell. A stimulus with the same orientation and strength was placed in the surround, and the response was recorded. Attention was then directed either to the center or surround stimulus. The mean responses relative to the center alone is shown for a sample of 100 neurons from the 2-D model. Error bars indicate the standard error of the mean. All three response groups are significantly different from each other at *p* < .005 (student’s t-test).

We simulated this experiment in the 2-D model as well. 100 neurons were randomly selected from the network. For each neuron, we measured the response to a strong stimulus of preferred orientation centered on the receptive field, and then added a strong stimulus of the same orientation to the surround. The response of the cell was measured in the absence of an attentional input (the “Attend Away” condition), as well as with an attentional input directed towards the center or surround stimulus. As was observed experimentally and in our line model, attending to the surround boosted the amount of surround suppression, whereas attending to the center greatly weakened the surround suppression (Figure 4B, compare the results of the 2-D model to the inset of Figure 4A).

### 2.3. Experimental paradigm alters the impact of attention

#### 2.3.1. Effect on contrast and response gain

All of the experiments and simulations discussed thus far demonstrate that attention produces a gain change in the firing rate of neurons within the locus of attention. The quality of this gain change, however, can be strongly influenced by the relative sizes of the stimulus and the attentional field. Reynolds and Heeger (2009) (their Figure 3) found in their normalization model of attention that when attention is directed to a relatively large area, the effect on the response to a small stimulus should be predominantly a change in “contrast-gain”, such that cells respond to stimuli as if they were effectively at higher contrast. This would be seen as a leftward shift in a contrast-response curve for a stimulus, with relatively little change in the maximum firing rate. For a large stimulus and a small attentional field, they instead predict a change in “response-gain”, such that all responses are scaled multiplicatively.

Here we again employ the one-dimensional spatial line network model to study the two different effects of attention described by Reynolds and Heeger (2009). Attention was still modeled as a small additional input only to excitatory cells over a defined spatial area, and we calculated “contrast response curves” with and without attention. (Note that what we call “contrast” is actually external input strength, *i.e*. the parameter *c* in Eq. 3; in reality, external input strength, as measured by thalamic input firing rate, is a monotonic but nonlinear, saturating function of stimulus contrast, (*e.g*. Sclar, 1987; Sclar et al., 1990).) To quantify changes in the contrast response properties, we fit each curve to a standard Naka-Rushton equation (Naka and Rushton, 1966):

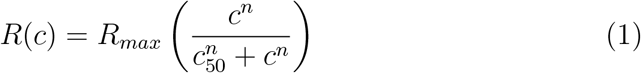

where *R_max_* is the plateau firing rate, *n* describes the steepness of the contrast response curve, and *c*_50_ is the strength of the stimulus at which the response is 50% of its maximum. In our fitting procedure, the value of n is discovered for the no-attention condition, and held at that value when fitting the attended condition.

With a large attentional field and small stimulus, the effect of attention was predominantly a leftward shift in the contrast-response function, as predicted by the model of Reynolds and Heeger (2009). We quantified this change in “contrast gain” as the difference in the *c*_50_ parameters of the contrast response curves produced with and without attention (Figure 5A, left). We compared this to the “response gain”, which we quantify as the ratio of *R_max_* parameters with and without attention. With a large stimulus and small attentional field, the effect of attention was reversed: there was little change in the contrast gain, and a much larger change in the response gain (Figure 5A, right). The dashed lines in either figure show the percent change in firing rate induced by attention. With a change in contrast gain there is little change in firing at the largest contrast, but this is not true for a change in response gain.

**Figure 5:**
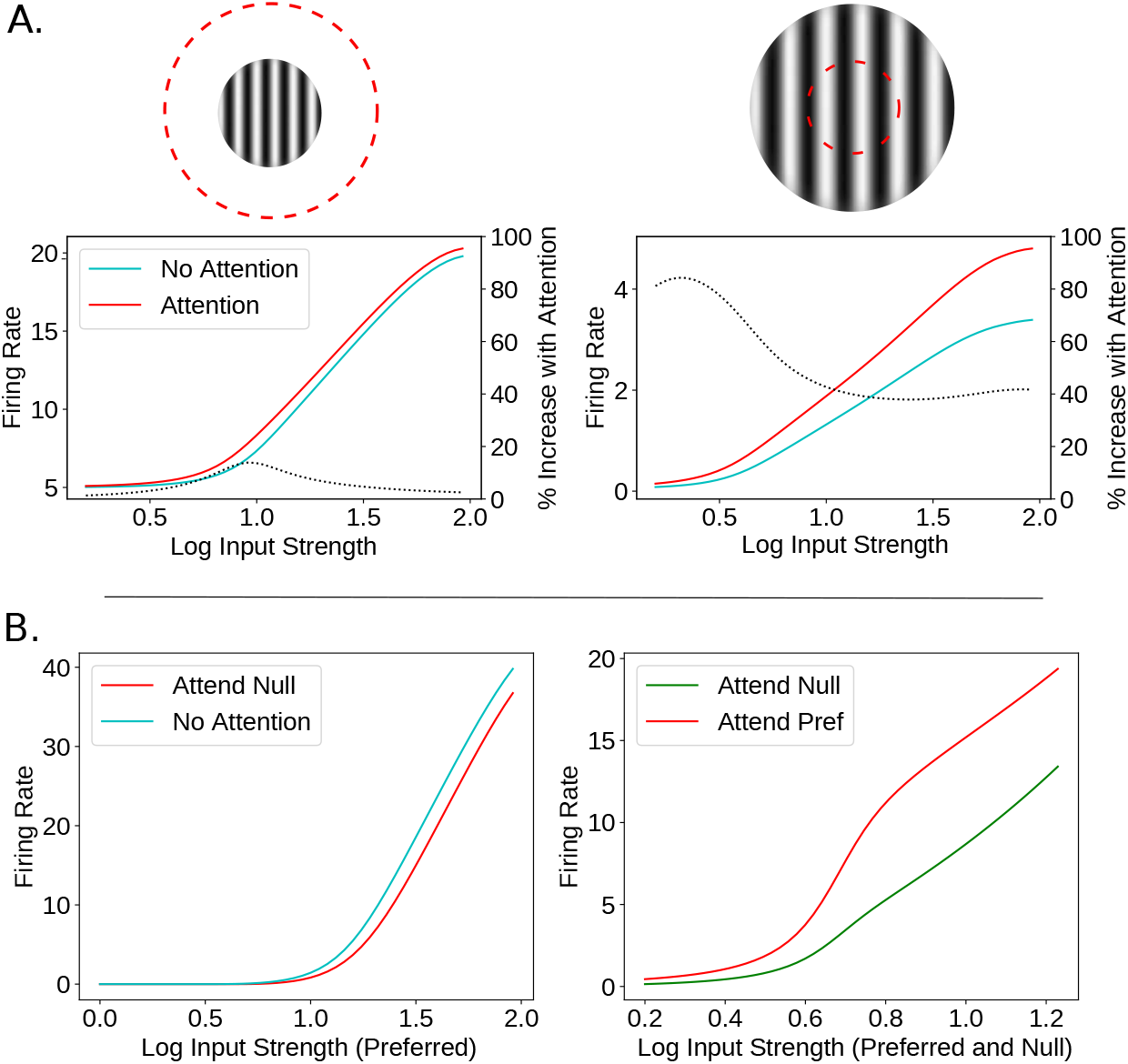
A.) The qualitative effect of attention depends on the relative sizes of the attentional and stimulus fields. Here we used the spatial line model to study the two different effects of attention, as described by Reynolds and Heeger (2009), Figure 3. Contrast response curves were calculated by varying the input strength logarithmically (base 10) in the presence (red curves) and absence (cyan curves) of attention. Left: with a large attentional field (red dashed circle) and small stimulus, the impact of attention was largely on contrast gain, defined as the difference between *c*_50_ values with and without attention (*R_max_* ratio: 0.98, *c*_50_ difference: −6.43). Right: in the “small attentional field, large stimulus” condition, attention mainly affected response gain, defined as the ratio of *R_max_* values (*R_max_* ratio: 1.39, *c*_50_ difference: −0.88). Dotted lines show the percent change in firing caused by attention. B.) Experimental paradigm alters gain change type. In the ring model, in the presence of a fixed-strength non-preferred stimulus (left), the contrast of a preferred stimulus was varied logarithmically (base 10) while attention was directed either away (cyan) or towards the non-preferred stimulus (red) as in Figure 4 of Reynolds and Heeger (2009). Attention to the non-preferred stimulus produced mainly a reduction in contrast gain, measured as the difference between *c*_50_ values (*R_max_* ratio: .97, *c*_50_ difference: 5.94) (Martinez-Trujillo and Treue, 2002). Showing preferred and non-preferred stimuli of equal but varying contrast (right) while attending to one or the other produced a much larger change in response gain, measured as the *R_max_* ratio (*R_max_* ratio: 1.38, *c*_50_ difference: −2.17). This was studied experimentally in Lee and Maunsell (2009).

While Reynolds and Heeger (2009) showed this property in their descriptive model of attention, conditions that produce changes in contrast or response gain have also been shown experimentally. Martinez-Trujillo and Treue (2002) recorded from neurons in area MT while presenting two stimuli within the receptive field. One stimulus was moving in a preferred direction, and the other in a non-preferred direction. They then varied the strength of the preferred stimulus while holding the contrast of the non-preferred stimulus fixed, and directed the monkey to attend either to the non-preferred stimulus or outside of the receptive field. They found that attending to the non-preferred stimulus caused predominantly a change in contrast-gain. However, Lee and Maunsell showed that if the contrast of both the preferred and non-preferred stimulus were varied simultaneously, attending to one or the other stimulus would produce a much larger change in response gain (Lee and Maunsell, 2009). Using the ring model again, we modeled both of these stimulus conditions, and find analogous results (Figure 5B).

#### 2.3.2. Effect on length tuning and receptive fields

Spatial attention has also been shown to change some of the basic features of a cell’s receptive field such as the size of the classical receptive field and the location of the center and surround.

The impact of spatial attention on length tuning was explored in Roberts et al. (2007). In this study, the length of an oriented bar was varied as firing rates from V1 cells were recorded. Attention was directed to the stimulus or to a stimulus in the opposite hemifield. The authors found that, for receptive fields near the fovea, attention had the effect of decreasing preferred length (that is, the length of the bar that elicits the highest firing rate). For receptive fields in the periphery, the reverse was true: attention increased the preferred length.

We explored attention’s impact on length tuning using the spatial line model. For different lengths of the stimulus, firing rates were recorded from a neuron at the center. The effect of attention varied as a function of the size of the attentional field. In Figure 6A (right) the ratio of the size of attention to the size of the stimulus is on the x-axis. By keeping a fixed ratio of attention size to stimulus size, we assume that the size of the attentional field scales with the size of the stimulus, but this scaling factor may differ for different cells. For small values of this attention scale factor, the preferred length with attention was greater than the preferred length without it. For higher values, this ratio was reversed. On the left, the firing rate as a function of length for two different values of this attention scale factor are shown. This pattern of how attention impacts preferred lengths reflects the impact of attending to the suppressive surround. With attention larger than the stimulus, more of the suppressive surround is activated for any given stimulus length. This effectively increases the length of the stimulus, making the preferred length smaller than without attention.

**Figure 6:**
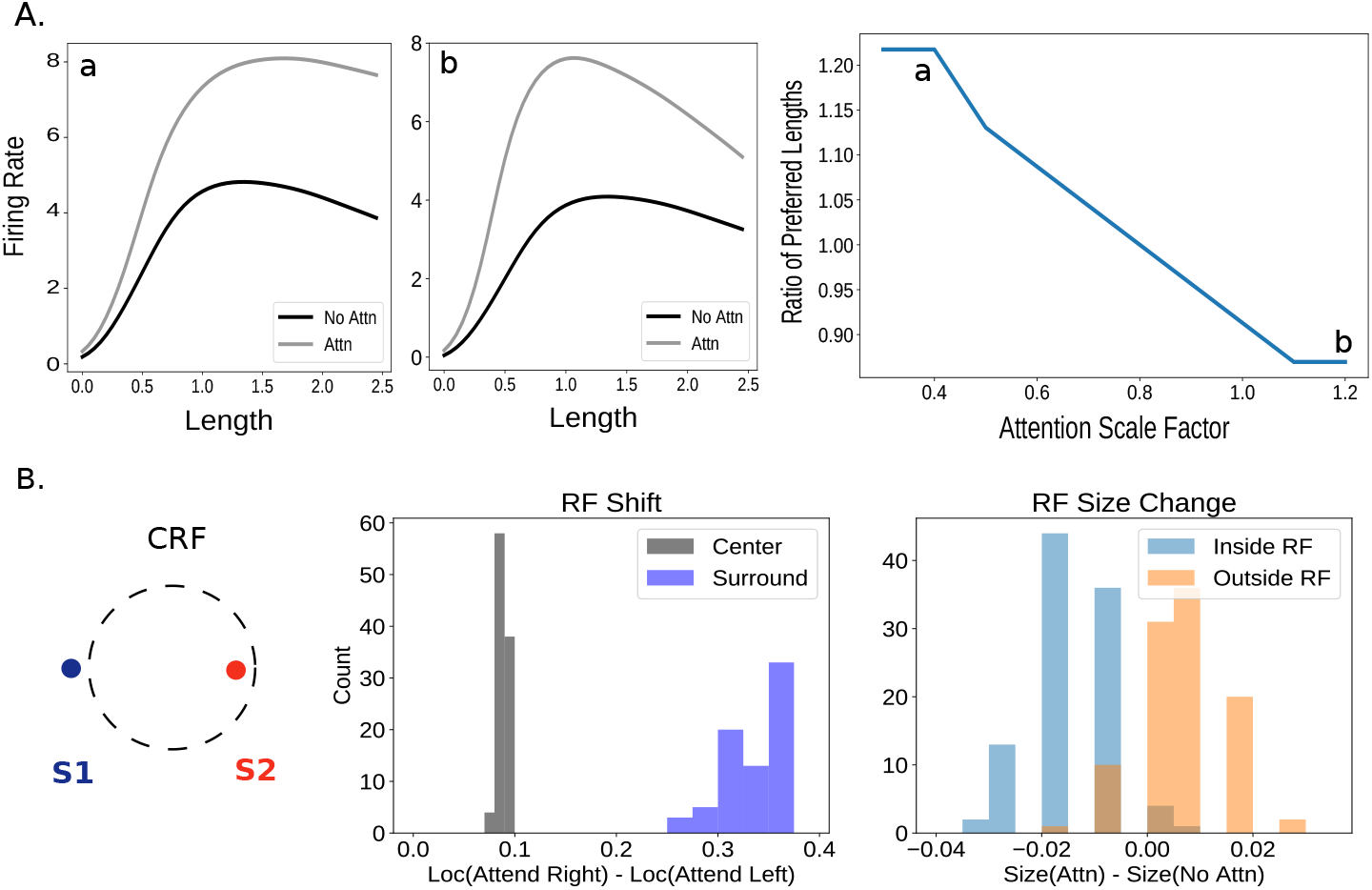
A.) Size of attention influences length tuning. Using the line model, we presented a stimulus of increasing length (left two plots). If attention was small compared to the stimulus (far left) attention shifted the preferred length (i.e., the length that elicits the highest firing rate) rightward, making it larger. If the area to which attention was applied was large compared to the stimulus (middle), the opposite occurred. Thus, varying the ratio of the size of attention to the stimulus size (“attention scale factor”) caused a shift in the ratio of the preferred lengths (preferred length with attention divided by preferred length without attention; right plot). Scale factor in the far left plot is marked on the right plot by the letter A, middle by B. In Roberts et al. (2007) the ratio of preferred lengths for parafoveal receptive fields was .88 and for peripheral receptive fields 1.19. B.) Shift and size change of receptive fields. The experimental setup involves two stimuli, the centers of which are placed on the edges of the classical receptive field (CRF) either slightly inside or outside (exact locations of stimuli chosen from a distribution). The locations of the receptive field center and surround were measured when attention was directed either toward the stimulus on the right edge or on the left edge. The difference in these locations between attentional conditions (normalized by cRF size) is shown in the left historgrams (positive values indicate rightward movement). The diameter of the cRF was also measured under these conditions and under a no-attention condition. The change in cRF size compared to the no-attention condition when the stimulus was slightly inside the cRF and slightly outside are shown in the right histograms (normalized by cRF size in the no-attention condition).

Our results combined with the findings of Roberts et al. (2007) suggest that attention targets parafoveal receptive fields differently than it targets peripheral ones. In particular, spatial attention inputs to parafoveal cells may be larger than the size of the stimuli these cells respond to. In the periphery, spatial attention inputs may represent an area smaller than the stimulus. This could be a result of the differently sized receptive fields in these two regions.

In Anton-Erxleben et al. (2009), the authors explored the impact of spatial attention on receptive field location and size. Two weak stimuli were placed near opposite edges of a cell’s classical receptive field (cRF). Attention was directed to one or the other, while a strong probe stimulus mapped the receptive field in these different conditions. We replicate this in the line model and find (Figure 6B) that the location of the receptive field center shifts toward the attended stimulus by an average of 8.9% of the cRF length. The center of mass of the surround shifts by an average of 34.8% in the same direction (the corresponding experimental values are 10.1% and 20.2%, respectively). We also measure how the size of the cRF changes with respect to a no-attention condition. We find that the cRF width shrinks when attention is deployed to a stimulus slightly inside the cRF (average size change of −1.5% of cRF area) whereas it grows when deployed to the stimulus slightly outside of the cRF (+.63%). The same qualitative relationship is found in Anton-Erxleben et al. (2009), though the magnitude was larger (−4.7% and + 14.2% respectively), and they reported a weak correlation between the location of the attended stimulus with respect to the cRF boundary and size change (*r* = .4).

Importantly, while our replication of the receptive field shift is robust to different parameter values and modeling details, the size change is both weaker and depends on certain specific choices. For example, if the stimuli are too weak, we only see shrinkage of the cRF width, and if the stimuli and probe are too small compared to the cRF size, the relationship between stimulus location and size change weakens. Most saliently, however, in our simulation, we always place one of the two stimuli slightly outside of the cRF and the other slightly inside (with the exact locations drawn from a distribution, see Methods 4.2). If we allow both to be inside or both outside, the relationship between the location of the stimulus and receptive field size change disappears. In Anton-Erxleben et al. (2009), the exact locations of the stimuli pair for each cell are not stated, but it is possible that by restricting their analysis to only cells that had one stimulus in and one out of the cRF they may have found a stronger relationship between location and receptive field size change.

Previous models have suggested that these receptive field changes may be due in part to feedforward effects from attention acting on the inputs to a circuit (Compte and Wang, 2006; Miconi and VanRullen, 2016). Such effects are not present in our one-layer model but may be important to incorporate in order to fully explain these findings.

#### 2.3.3. Factors influencing the magnitude of attentional effects

In Lee and Maunsell (2010), the authors controlled attention and task difficulty across stimulus conditions while varying the number of stimuli in the receptive field of MT neurons. Through this, they showed that attentional modulation was weaker when only one stimulus was present in the receptive field, and that this result is well-captured by a divisive normalization model. We use the ring model to replicate these results. By presenting three different stimuli (a most-, moderately-, and least-preferred orientation) either alone or in pairs (Figure 7A, left; compare to Lee and Maunsell (2010) Figure 4), we show that the effect of an attentional input was strongest when applied to one stimulus in a pair. In particular, effects of attention on firing rates were highest when moving attention from outside the receptive field to the preferred stimulus inside the receptive field when a non-preferred stimulus is also present (Figure 7A, right). The next strongest effect was from moving attention from the non-preferred stimulus in the receptive field to the preferred. Finally, attention effects were weakest when moving attention from outside the receptive field to a preferred stimulus presented alone inside the receptive field.

**Figure 7:**
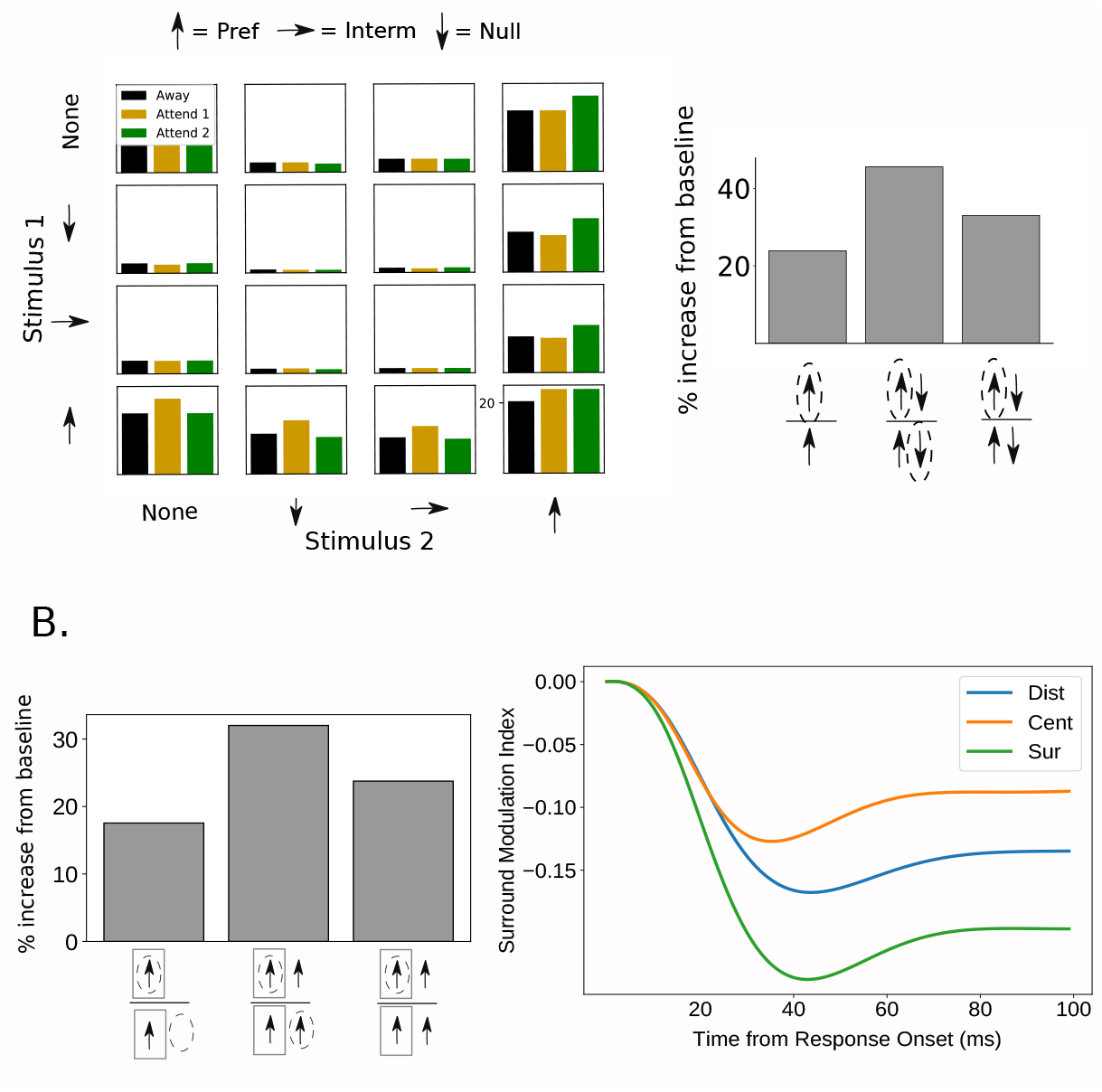
A.) Effects of attention are greater with more than one stimulus in the receptive field. Using the ring model, three different stimuli (preferred, intermediate, and null) were shown either individually or in pairs. Attention was directed to either of the two stimuli (‘Attend 1’ or ‘Attend 2’) or outside of the receptive field (‘Away’; when only one stimulus was present, attending to the opposite stimulus is the same as attending away). Left: Bar plots represent steady state firing of the recorded neuron for all stimulus and attention conditions. Right: bar plots indicate percent increase in firing rate with attention, for three different comparisons. Arrows indicate which stimuli were in the receptive field for the two conditions being compared (bottom arrows indicate baseline condition, top arrow(s) indicate attended condition) and dashed circles indicate attended stimulus. The comparable values for these conditions from Lee and Maunsell (2010) are 9%, 59%, 28% respectively. B.) Effects of attention are greater with a stimulus in the surround. Using the line model, a preferred stimulus was presented in the receptive field center. Left: bar plot indicates increase in firing in preferred-attended condition (top arrows) vs. baseline condition (bottom arrows). Rectangles indicate receptive field. The presence of a surround stimulus is indicated by an additional arrow outside the receptive field and attention is indicated by a dashed circle. The increase in firing was smaller without the surround present (comparable values from Sundberg et al. (2009) are 18.8% versus 36.8%. The authors do not report the percent increase compared to a baseline condition without attention to either center or surround). Right: the strength of firing rate modulation from the addition of a surround stimulus (the surround modulation index: [*r*(*C* + *S*) – *r*(*C*)]/[*r*(*C* + *S*) + *r*(*C*)]) is plotted vs. time, for different attention conditions: attending the surround, attending the center, and attending a distant location (modeled as no attention). The difference between these conditions emerged over time.

A similar comparison was done using spatial attention rather than feature attention in Sundberg et al. (2009). Here, attention was moved between the receptive field center and the suppressive surround. A stimulus of preferred orientation was present in the center and was present or absent in the surround. The impact of attending the center was larger when the stimulus in the surround was present (Figure 2 of Sundberg et al. (2009)). We replicated these results using the line model. The firing rate of an excitatory cell was recorded with a stimulus centered on its preferred location. Attention was applied to this location, or to a location in the surround both in the presence and absence of a stimulus there. The results of this are shown in Figure 7B (left).

In Sundberg et al. (2009), the impact of attention on surround suppression was also shown over time. The extent to which firing rates are decreased by the presence of the surround was measured when attention was directed to the receptive field center, surround, or to a distant location. The authors note (their Figure 5) that the difference in surround modulation between these different attention conditions emerged over time. Our model shows the same result (Figure 7B, right). This demonstrates how our circuit-level implementation of the normalization model of attention goes beyond just capturing steady state effects. It can also replicate dynamics.

Note, that the differences emerge faster in our model than in the data (in the data, the difference is not seen in the time bin 15-55ms after response onset, but emerges sometime in the next 40ms time bin). However, our model does not take into account any delays in the onset of the attentional signal relative to the onset of stimulus-driven feedforward input to the recorded neurons.

### 2.4. Attention reduces trial-to-trial variability and noise correlations

In addition to its effects on mean firing rates, attention has also been shown to modulate the variability in rates across trials. Mitchell et al. (2007) showed that attending to a stimulus decreased the across-trial variability of neural responses when compared to trials in which attention was directed elsewhere. Furthermore, this experiment showed that this decrease in variability occurs in both broad spiking (putative excitatory) cells and narrow spiking (putative inhibitory) cells.

To study this effect in our model, we introduced a source of trial-to-trial variability into our ring network by given each neuron a noisy input in addition to its stimulus inputs, similarly to Hennequin et al. (2018) (see Methods 4.1.1 for details). We then ran 1,000 trials of a simple stimulus presentation. On half of these trials, attention was directed towards the stimulus being presented. On the other half there was no attentional modulation added to the network. The stimulus onset produced a reduction in the trial-to-trial variability, measured as the Fano factor, with this reduction occurring both for neurons that are activated by the stimulus and neurons that are not activated or suppressed (Figure 8A), as in experiments (Churchland et al., 2010) and as previously shown for the SSN (Hennequin et al., 2018). Addition of attention caused an additional drop in Fano factor, again regardless of whether the stimulus plus attention caused a net increase, zero change, or net decrease in firing rate (Figure 8A, right).

**Figure 8:**
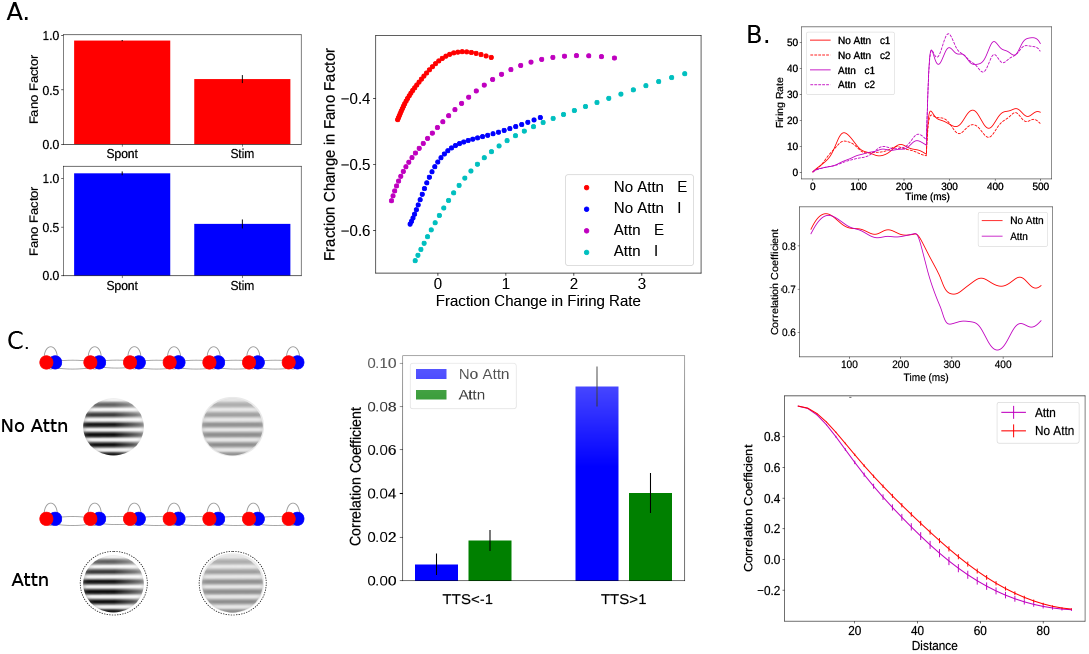
A.) Attention causes a reduction in trial-to-trial variability. In the ring model with noisy background input, 35 E (red) and 35 I (blue) cells were recorded as a stimulus that was oblique (but not orthogonal) to their preferred stimuli was presented. Stimulus onset produced a substantial reduction in trial-to-trial variability, measured as the Fano factor, compared to spontaneous activity (left; errorbars are STD). On the right, fractional change in Fano factor is plotted as a function of fractional change in firing rate for each of the 35 E and 35 I cells in the presence and absence of attention. In all cells, stimulus onset produced a decrease in the trial-to-trial variability, regardless of whether the stimulus produced an increase, decrease, or no change in the mean firing rate (Churchland et al., 2010). In the presence of attention, this decrease in variability was enhanced. The percent change in both firing rate and Fano factor was calculated for each cell by taking a time average of both the mean rate and Fano factor before and after the onset of the stimulus (in trials with attention, it came on at the same time as the stimulus). B.) Attention decreases noise correlations between neurons. In the ring model with noisy background input, stimulus onset produced a reduction in noise correlations between pairs of neurons in the network. The mean firing rates of two nearby excitatory cells (c1, c2) in each of the two conditions is plotted on top; stimulus (at 90 degrees) and attention turn on at 250ms. The correlations between the two cells are plotted below it. Correlation time-series are shown as a running average with a 50-ms sliding window. On the bottom, the mean correlation between pairs of recorded cells (representing 30-65 degrees) during the stimulus response epoch is plotted against difference in preferred orientation. Error bars indicate SEM. C.) Attention increases or decreases noise correlations between neurons based on preferred stimulus. In Ruff and Cohen (2014), animals performed a contrast discrimination task on two nearby stimuli, represented here as two inputs to the line model of different strengths. During different blocks, attention was directed to one of two such sets of stimuli, one in each hemifield. Here we model attention to the opposite hemifield as a ‘no attention’ condition (top left) and attention to the hemifield of the recorded cells as attention to each of the two stimuli simultaneously (bottom left). The 25 model cells we analyzed responded to one or the other stimulus alone. TTS values are the product of d-primes and represent whether a pair of cells has the same (positive) or different stimulus preference (negative). By creating 20 populations of 25 cells each, we analyzed the relationship between TTS and the effect of correlation on attention for 6000 cell pairs. Through this we found both a significant (*p* << .05) decrease in correlation with attention for cells that preferred the same stimulus and increase for cells that had opposite preferences (right). Error bars indicate SEM. For more details, see Methods 4.2.

In addition to causing a drop in trial-to-trial variability, Cohen and colleagues demonstrated that an even stronger effect of attention on network variability is a pronounced decrease in the magnitude of noise correlations between neurons in V4 (Cohen and Maunsell, 2009). This aligns with the finding that a stimulus suppresses the shared or correlated component of neural variability, not the component private to each neuron (Churchland et al., 2010). Cohen *et al*., 2009, recorded from thousands of pairs of neurons and multiunit clusters in V4 during a visual change detection task, and found that the presence of attention greatly enhanced performance. They further argued that the significant improvement in performance was not due to changes in single neurons, but rather to a pronounced drop in noise correlations.

To simulate this experiment, we recorded from pairs of excitatory cells in the ring model in the presence of noisy input while presenting the network with two high-contrast oblique stimuli. On half of the trials, attention was directed to one of the stimuli. We calculated the correlation between all pairs of recorded neurons in the presence and absence of attention. Pairs of neurons were grouped based on their distance from each other on the ring (i.e. difference in preferred orientation). The changes in firing for two example neurons with attention as well as the noise correlations between them over the course of an example trial are shown in Figure 8B (top). The average value of noise correlations between neurons at various distances is shown on the bottom. As was observed experimentally, attention caused a reduction in the noise correlations between neurons beyond the reduction caused by the stimulus alone.

The suppression of correlated variability can be understood as resulting from the normalization performed by the model (although the model also explains further aspects of this suppression not explained simply by normalization, Hennequin et al. (2018)). In particular, as has been observed experimentally (Busse et al., 2009), normalization averages the responses to approximately equal strength inputs but performs a more unequal averaging of unequal strength stimuli, becoming “winner-take-all” when inputs differ sufficiently in strength (Rubin et al., 2015). The reduction in correlated variability with increasing stimulus strength can be understood to occur because the ongoing noisy inputs become steadily weaker relative to the stimulus. The normalization thus increasingly favors the response to the stimulus and suppresses the noise. Because this suppression is mediated by the network, it acts on the correlated component of the noise and not on the private noise, which is largely averaged out in its impact at the network level.

An alternative picture of the mechanism of suppression is that it occurs through the enhancement of the strength of feedback inhibition with increasing network activation (Hennequin et al., 2018). In particular, in linearizations about the deterministic fixed point, the real parts of the leading eigenvalues become more negative with increasing mean stimulus drive, representing increased feedback inhibition of the corresponding eigenvector activity patterns onto themselves, dampening their fluctuations. Given structured connectivity, these activity patterns have similar structure and so their fluctuations represent correlated variability.

Investigations regarding noise correlations have indicated that a decrease in correlation with attention should only occur for pairs of neurons that represent the same stimulus whereas pairs of neurons representing different spatial locations or features may actually see an increase in correlations (Averbeck et al., 2006).

This bi-directional effect of attention was observed during a spatial attention task in cells in area V4 (Ruff and Cohen, 2014). In this task, subjects were required to perform a contrast discrimination task in the cued hemifield. To replicate this study we used the line model with two nearby stimuli of unequal contrast (Figure 8C, left). The TTS metric from Ruff and Cohen (2014) measures the extent to which a pair of cells have the same (positive TTS) or opposite (negative TTS) preferred stimulus of the two presented. Replicating Figure 5 from that paper, we see that attention decreased correlations for cells with the same preferred stimulus but increased it for those with opposite preferred stimuli (Figure 8C, right).

As can be seen in Figure 8B, in the simulation we ran on our ring model, attention decreased correlations for cell pairs with a wide range of differences in preferred features. To be analogous to the results of Ruff and Cohen (2014), cell pairs with large differences in preferred features should show an increase in correlations with attention. This result did occasionally occur in our ring model when using weaker stimuli and/or a smaller number of trials to calculate the correlations in the ring model. Examples of this can be found in Supplementary Figure A.11.

Reasons why the bi-directional change in correlations is seen with spatial attention in our line model but not (consistently) with feature attention in our ring model may have to do with the reach of inhibition in these two models. In our line model, inhibitory populations project only to the E and I populations at the same spatial location; in the ring model, however, inhibitory connections are broader, projecting to E and I cells with different preferred features. If we alter our line model to have broader inhibitory connections, it fails to replicate the findings of Ruff and Cohen (2014) (not shown). It would be interesting to see if this difference between spatial and feature attention can be seen experimentally.

### 2.5. An alternative mechanism

In all of the simulation results presented thus far, attentional modulation has been modeled as a small excitatory input biased towards the excitatory cells within the locus of attention. Here we consider instead a small inhibitory input to inhibitory cells within the locus of attention, disinhibiting rather than exciting the excitatory cells. This is motivated by two observations. First, it was observed that inputs from Anterior Cingulate Cortex to V1 target the VIP class of inhibitory cells (Zhang et al., 2014). The VIP cells in turn are known to inhibit other inhibitory neurons and, at least in V1, disinhibit excitatory cells (*e.g*. Fu et al., 2014). The ACC input conceivably could be involved in attentional modulation. Second, electrophysiologic work has revealed the function of two classes of inhibitory cells in layer 1 of cortex (Jiang et al., 2013). One of these classes, the single bouquet cells (SBCs) was shown to preferentially inhibit the interneurons of deeper layers, and so have a net disinhibitory effect on the local pyramidal cells. As layer 1 receives a significant portion of its input from higher cortical areas, it has been suggested that this circuit may play a role in attention and other top-down modulation of local circuit activity (Larkum, 2013).

To test the feasibility of this mechanism in our model, we repeated our suite of simulations using this alternative, disinhibitory mechanism of attention. Rather than modeling attention as an additional excitatory input to E cells, we instead model it as an additional inhibitory input to I cells. The results of these simulations are presented in the Supplementary Figures. Overall, this alternative mechanism can qualitatively reproduce most of the findings we report above (Supplementary Figure A.12). Frequently, however, the same value of the attention strength parameter produces weaker effects on neural firing than when attention is directed towards the excitatory cells (for example, compare Figure 7B to Figure A.12G).

In addition, there are instances where this form of attention does not qualitatively replicate our original findings (Figure 9). One major discrepancy between results comes from the use of the 2-D model. Comparing Figure 9B to Figure 3, modeling attention as inhibition to inhibitory cells creates the opposite relationship (i.e., a negative correlation) between attentional modulation and normalization. Note also that the AMI’s in Fig. 9B are small and of both signs, meaning that attention to the preferred stimulus does not consistently cause greater increases in firing rate than attention to the anti-preferred stimulus under this model of attention. In the 2-D model, any additional inhibitory input to the inhibitory population has the effect of increasing firing rates for many of the cells, even those representing unattended stimuli. The model therefore cannot replicate findings that rely on attention to a non-preferred stimulus causing a decrease in firing rate. This appears to be a consequence of the strong inhibition needed to keep this more complex model in a stable regime. Attention directed toward inhibitory cells also has a surprising effect on the correlations explored in Figure 8C. As can be seen in Figure 9E, this form of attention increases correlations for pairs of cells both with the same and opposite preferred stimuli.

**Figure 9:**
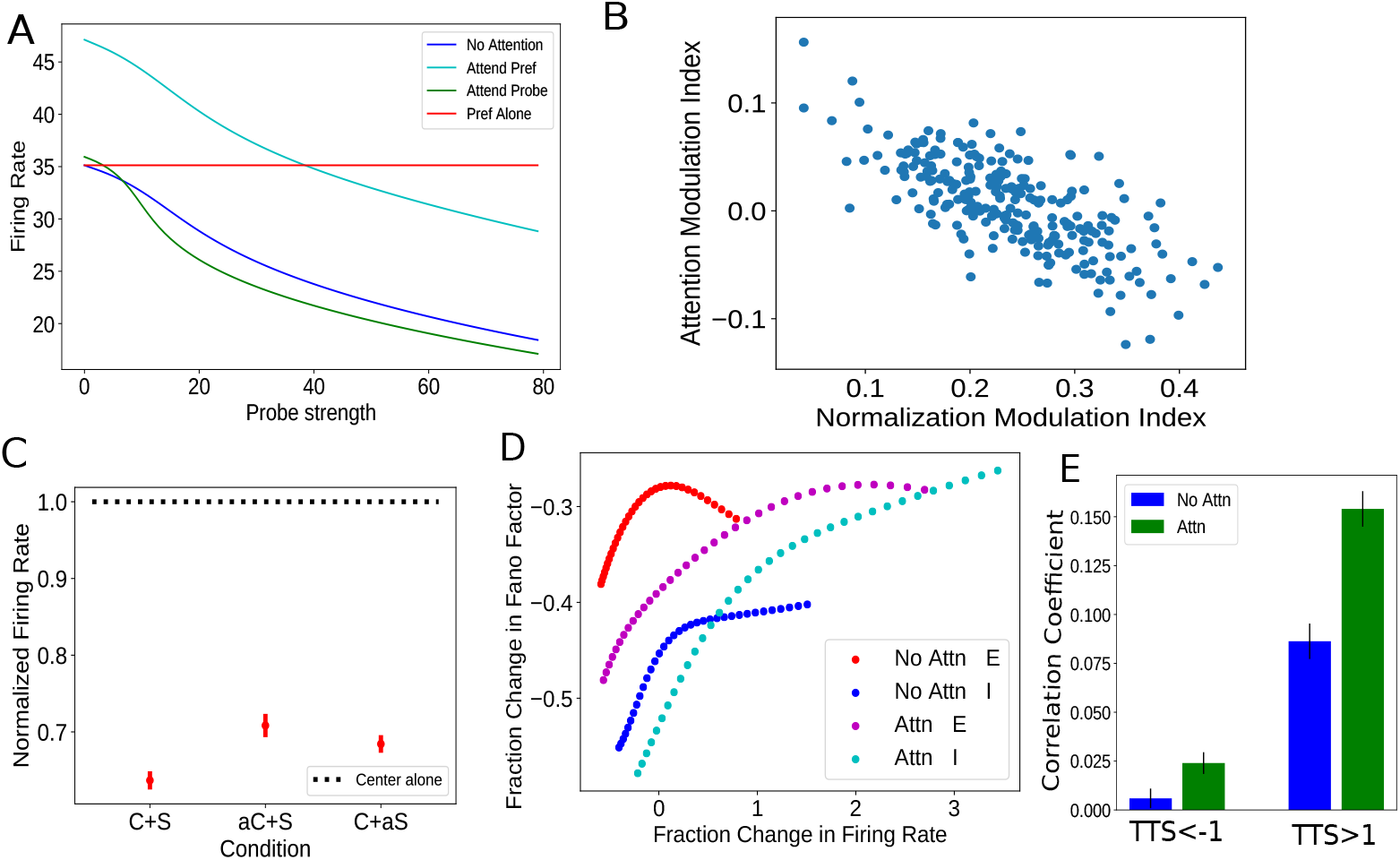
**Findings not qualitatively replicated with attention modeled as inhibitory input to inhibitory cells** A. Figure 2A. Here most of the results are replicated, however at low probe strengths attending the probe can increase firing rates compared to no attention. B. Figure 3. Here the relationship between normalization and attention is negative. C. Figure 4B. Here the attend-surround condition is too similar to the attend-center one. D. Figure 8A. Here for a range of firing rate changes, inhibitory cells have their Fano Factor increased with attention (though it should be noted this result happens occasionally when modeling attention as excitation to excitatory cells, for example, when the number of trials is lower). E. Figure 8C. Here cell pairs with TTS>1 also show an increase in correlation with attention.

### 2.6. Attention enhances detection performance in a multi-layer model

An important consequence of deploying attention is enhanced performance on challenging tasks. We have thus far shown how the SSN can replicate many neural effects of attention, but to truly understand attention, it is necessary to link these neural changes to performance changes. And for that it is necessary to build a functioning model of the visual system that can perform visual tasks.

Because the SSN replicates neural findings that have been found in various areas in the visual system, it can be thought of as a canonical circuit, which is repeated throughout the visual hierarchy. To build a biologically-realistic multi-area model of the visual system that can perform a task, we therefore model each area as a set of SSNs, the outputs of which are fed into another set of SSNs (i.e., a downstream visual area). The precise connections between these areas are learned as part of a training procedure.

In particular, the SSN circuitry is placed inside a convolutional neural network architecture, creating a model we have dubbed the SSN-CNN (Methods 4.3). The structure of the model can be seen in Figure 10A. The network is a 2-layer convolutional neural network wherein the convolutional filters are constrained to be non-negative (to mimic the excitatory feedforward connections that exist between different visual areas). In addition, after each pooling layer is an SSN layer. The SSN layer implements normalization dynamically (historically normalization layers have been included in CNNs, typically implemented via a divisive normalization equation Krizhevsky et al. (2012)). Specifically, at each spatial location on the 2-D map, a ring SSN implements feature normalization across the different feature maps. During training, the recurrent connections of the SSN layers are held constant while all other weights of the network are trained end-to-end via backpropagation through time on the MNIST 10-way digit classification task.

**Figure 10:**
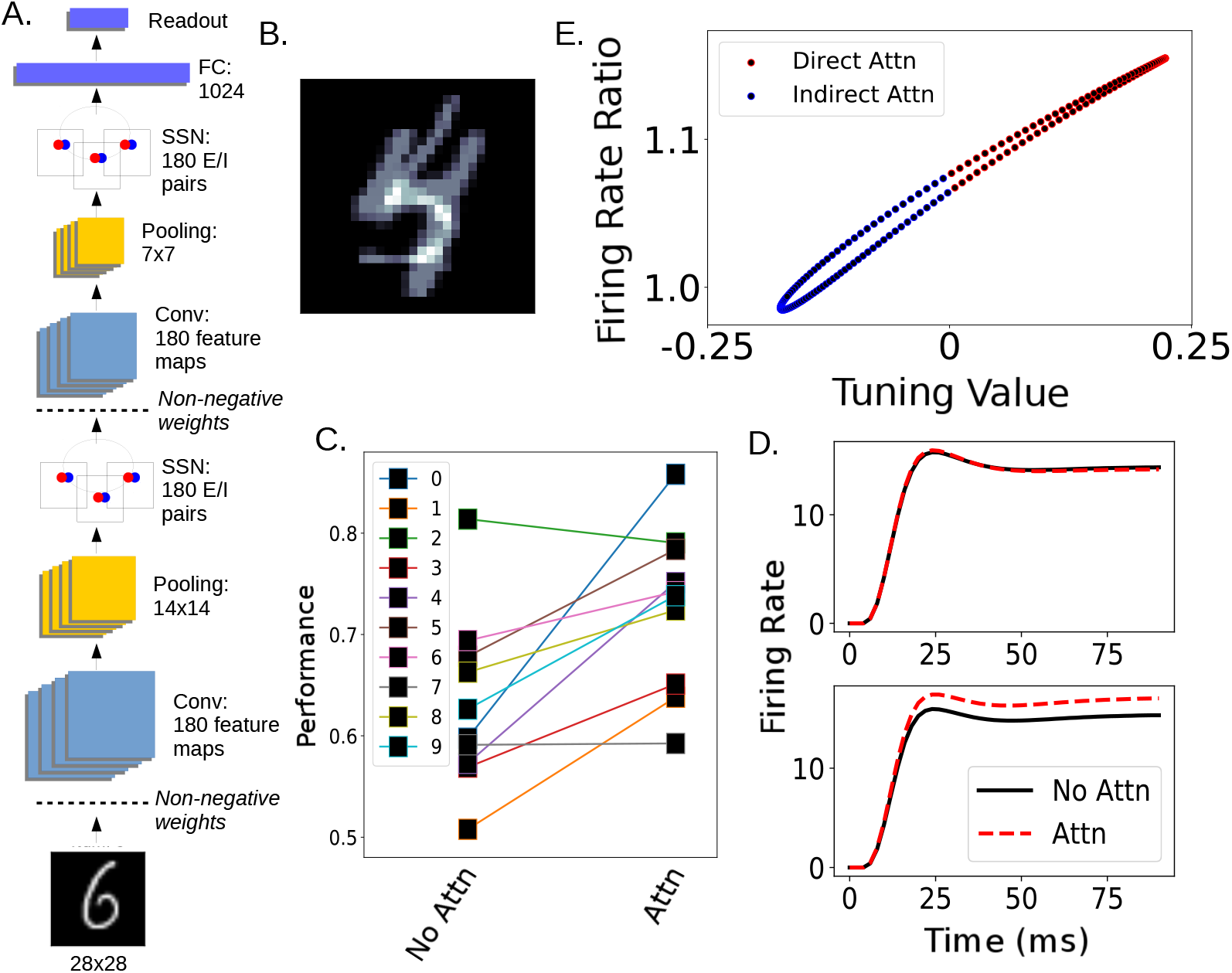
Attention in the SSN-CNN enhances visual detection performance. A.) The architecture of the SSN-CNN model. In the SSN layers, a full ring model exists at each spatial location (though only one is shown). B.) An example of the images used in the attention task. This image contains a ‘5’ and ‘4’ overlaid, therefore both the binary classifier trained to detect 4s and the one trained to detect 5s should respond positively. C.) Binary detection performance for each digit with (right) and without (left) attention. D.) Example firing rate of two neurons recorded from the second SSN layer with receptive fields at the center of the image when shown the image in (B). The top neuron had a small decrease in firing when attention was deployed to the digit 4 and the bottom had an increase. E.) Impact of attention to the digit 4 on firing rates of excitatory cells (rate with attention divided by rate without) as a function of tuning to the digit. A feature map’s tuning value for a given digit is defined as its z-scored mean response to that digit (see Methods, section 4.3). Attention is modeled as excitatory input applied to feature maps whose tuning value is above the median value across maps for that digit. The strength of a map’s attentional input is proportional to the difference between that map’s tuning value and the median value. Only neurons marked in red were above the median and given direct attentional input.

Replicating Lindsay and Miller (2018), after the network is trained on the standard task, the final layer is replaced by a series of binary classifiers, one for each digit. These binary classifiers are trained on digit images to determine if a given digit is present in the image or not (for example, one of the binary classifiers would be trained to classify images as being of the digit ‘4’ or not). To test the impact of attention on the abilities of these binary classifiers, we presented the network with a more challenging task: determining if a given digit is present in an image that contains two overlaid digits (Figure 10B). The network performs above chance on this challenging task in the absence of attention.

Attention is applied in this model as previously described: an additional positive input is given to excitatory cells that prefer the attended digit. To determine which cells in the SSN layers “prefer” the attended digit we created tuning curves based on the response of excitatory cells in the SSN when presented with images of different digits (See Methods 4.3). Applying attention in this way still elicits attentional changes in the cells that are not directly targeted—through the recurrent connections—as can be seen in Figure 10E. This includes decreasing the firing rates of neurons that do not prefer the attended digit. While this feature attention is applied the same way across all the ring networks representing different locations in a layer, the pattern of feedforward input will influence the ultimate impact of attention. This can be seen by comparing the ratio of firing with and without attention in ring networks at different nearby spatial locations, which receive slightly different feedforward input (Supplementary Figure A.13).

Applying attention in this way at layer 2 of this network enhanced performance on the overlaid digit detection task (Figure 10C). This shows that the neural changes implemented by the SSN can cause performance changes.

Previous work (Lindsay (2015); Lindsay and Miller (2018)) has shown how attentional changes in different layers of a standard deep convolutional neural network can lead to enhanced performance on challenging visual tasks. That work demonstrated that the attentional modulation style that works best is multiplicative and bi-directional changes (i.e., the effect of attention should be to scale the activity of neurons that prefer the attended stimulus up and those that don’t prefer it down). What we have shown here is how an additive input solely to the excitatory neurons that prefer the attended stimulus can turn into multiplicative and bi-directional changes via the circuit mechanisms of the SSN and lead to an increase in performance. This allows for a straightforward mechanism by which top-down attentional signals can lead to enhanced performance simply by providing additional synaptic inputs to the right set of excitatory cells.

## 3. Discussion

We have shown that, given a simple model circuit, modeling attention as an additional input to excitatory cells representing the attended stimulus suffices to reproduce a wide variety of experimental results on attention in visual cortex (Anton-Erxleben et al., 2009; Cohen and Maunsell, 2009; Lee and Maunsell, 2009; Martinez-Trujillo and Treue, 2002; Mitchell et al., 2007; Ni et al., 2012; Reynolds and Desimone, 2003; Ruff and Cohen, 2014; Sundberg et al., 2009; Treue and Martinez Trujillo, 1999). The model circuit, the stabilized supralinear network (SSN), involves only a few assumptions: a supralinear (expansive) input/output function, and connectivity between cells that decreases in strength with increasing difference in retinotopic position or preferred features. This model circuit has been shown previously to replicate a wide set of nonlinear neural response properties in multiple visual cortical areas (Adesnik, 2017; Ahmadian et al., 2013; Hennequin et al., 2018; Liu et al., 2018; Rubin et al., 2015), including those often summarized as “normalization” (Carandini and Heeger, 2012).

Through the combination of balanced amplification (Murphy and Miller, 2009) and the nonlinearity of the SSN model, a small additional excitatory input to excitatory cells causes a nonlinear scaling of firing rates in a manner consistent with a number of experimental observations. Recurrent connections implement interactions between features and spatial locations. These simple elements are sufficient to account for variation, under different experimental paradigms, in the effects of attention on stimulus interactions, receptive field properties, gain changes, and variability and correlations, as well as in the magnitude of attention’s effects. We have thus not just shown how the normalization model of attention could be implemented by a neural circuit, but have also shown how changes not normally attributed to the normalization model (e.g., effects on variability and correlations) can be explained by this same circuit as well. We are not aware of any previous model that has attempted to replicate so many effects of attention simultaneously.

The ability to replicate all these effects via a small additional input to a subset of neurons provides a simple, plausible mechanism through which higher cortical feedback can implement attention. Previous work has identified areas in the frontal cortex that may be considered the source of top-down selective visual attention (Bichot et al., 2015; Paneri and Gregoriou, 2017; Zhang et al., 2014). Exactly how connections from these areas target visual areas to create the changes seen with attention is unknown. Studying these feedback connections can be challenging, as it requires detailed anatomical and physiological investigations across multiple brain areas. For this reason, narrowing the hypothesis space by identifying which mechanisms of feedback control are theoretically capable of implementing the known effects of attention is important. Here, we show that positive additive input to the excitatory neurons that prefer the attended stimulus can recreate the multiplicative changes observed in both E and I cells and both in cells that prefer and do not prefer the attended stimulus. Adding negative input to the inhibitory cells that prefer the attended stimulus can also replicate many of these effects. However, this fails to reproduce certain results, notably the positive correlation between modulation by normalization and by attention and the reduction in noise correlations between cells with similar stimulus preferences (Figure 9). We do not know if this is a fundamental problem with a disinhibitory model of attention or if it could be fixed by altering model connectivity.

Overall, these results show that feedback connections do not need to be directly responsible for all of the neural effects of attention. Instead, they only need to target a subset of neurons in a simple specific way and the local recurrent circuitry can take care of the rest.

In addition to replicating known findings, the set of models presented here can serve as test beds for future work on attention. In particular, experimental designs can be explored and precise predictions made before carrying out further experiments. Within this work we have discussed possible predictions relating to how attention targets parafoveal versus peripheral receptive fields, the impact of experimental design on receptive field shrinkage, and differences between spatial and feature attention in terms of their impact on correlations. There are also further findings from the experimental literature that could be explored in this model. For example, a two-layer version of the line model could serve as a means of exploring how attention decreases correlations within a visual area but increases them across visual areas (Ruff and Cohen, 2016).

Circuit models in neuroscience are frequently built to replicate and understand the relationship between anatomy and neural activity. Traditionally, these models do not perform a perceptual or cognitive task. Yet, an ultimate understanding of the circuitry of visual perception will need to replicate behavioral as well as neural findings. We work towards this goal here by incorporating the SSN model into a convolutional neural network that can perform digit recognition (the SSN-CNN). Through this, we connected the neural changes our model replicates to enhanced detection performance. This model also sets a precedent for how traditional approaches from computational neuroscience can be incorporated with the increasingly popular approach of using deep neural networks to study the brain (Kell and McDermott, 2019; Lindsay, 2020; Yamins and DiCarlo, 2016).

Further connections between neural changes and performance remain to be explored, and the SSN-CNN could be useful in this pursuit. For example, we did not incorporate noise into the SSN-CNN in this work. Using the noisy version of the ring model would allow for an exploration of how noise and correlation changes impact performance. We also did not attempt to model or replicate effects of attention on reaction time, which could be addressed using the model’s dynamics. Using the full 2-D model (instead of ring models at each spatial location) would also allow for an exploration of the effects of spatial attention and the interaction between spatial and feature attention.

Altogether, our modeling results show that a wide range of experimental findings concerning both neural and performance effects of attention can be captured with a single, unified model. This demonstrates how a simple top-down signal could be responsible for these effects, providing insight on the necessary inter-area connections for attention. We believe the model provides a basis for many further explorations of the mechanisms of attention.

## 4. Methods

Code will be publicly available upon publication.

### 4.1. Basic Circuit Models

In this study we employ several different configurations of a basic SSN circuit model, the central unit in all being an interconnected pair of excitatory (E) and inhibitory (I) cells. The two core models are the one-dimensional ring model and the one-dimensional line model. In addition, for Figure 1A we use a simplified 2-cell circuit model, and for Figures 3 and 4B we use a large two-dimensional model.

In all models, each neuron, *i*, is represented as a firing rate unit whose activity, *r_i_*, evolves according to:

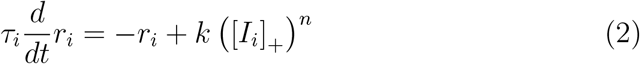

with *n* > 1 (indicating a supralinear activation function). The expression [*υ*]_+_ = max(*υ*, 0), that is, neuronal activity cannot go below zero. The inputs, *I_i_*, to a given neuron *i* are comprised of recurrent inputs, feedforward stimulus inputs, and attentional inputs. These inputs and parameter values are specified for each model below. In all models the time constant *τ_i_* has the value *τ_E_* = 20 ms for all E cells and *τ_I_* = 10 ms for all I cells. Simulations are run using the forward Euler method with time step 1ms.

All of these models except the E-I pair model were described previously in (Rubin et al., 2015), however we will recap them briefly here. In order to avoid results that depend on parameter tuning, we simply used the same model parameters as in that study (which did not address effects of attention), without tuning them in any way to get the current results. The models are only modified by the addition of attentional inputs, and by the addition of noise inputs for Figures 8A and B.

#### 4.1.1. Ring Model

The ring model is intended to represent neurons with a shared retinotopic receptive field but different preferred features. In this model, an E-I pair exists at each location on the ring, with the preferred feature (e.g. orientation or direction) varying smoothly around the ring. The relative input to a cell with preferred orientation *θ* from a stimulus of orientation *ϕ* is given by 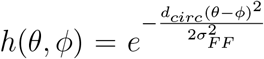 where *d_circ_*(*θ − ϕ*) is the shortest distance around the circle between *θ* and *ϕ*. The absolute stimulus input to a cell comes from multiplying *h*(*θ,ϕ*) by the scalar *c*, which represents the overall strength or contrast of the stimulus. In addition, attention directed towards orientation *ϕ*′ provides extra input to E cells with the same overall shape as a stimulus input, scaled by the attention strength factor, *a*. (In studies that modeled attention as negative input to I cells rather than positive input to E cells, this input is instead given to inhibitory cells, with *a* < 0.) In total, input to the E or I cell at location θ on the ring is given by:

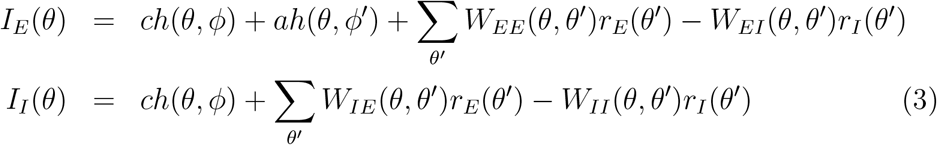

respectively.

Recurrent connections fall off according to 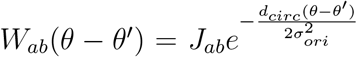, where *d_circ_*(*θ − θ*′) is the shortest distance around the circle between *θ* and *θ′*. If multiple stimuli are present the inputs are added linearly.

For simulations of this model, the following parameters are used: the number of E/I pairs is *N* = 180; the spacing in degrees between adjacent pairs on the ring is Δ*θ* = 1°; *J_EE_* = 0.044, *J_IE_* = 0.042, *J_EI_* = 0.023, *J_II_* = 0.018, *σ_ori_* = 32°, *σ_FF_* = 30°, *k* = 0.04, *n* = 2.0.

The ring and its inputs are schematized in Figure 1B.

In certain simulations, noise is added to the inputs to these cells. Specifically, 10 + *ν*(*θ, t*) was added to input to each unit at each timestep. External noise *ν* was given by convolution of unit-integral Gaussian temporal filter (stdev 10 ms) and spatial filter (stdev 8°) with Gaussian spatiotemporally white noise (mean 0, stdev 40), yielding 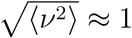.

#### 4.1.2. Line Model

In the line model, each E-I pair represents a different retinotopic location but all have the same preferred features. Rather than being arranged in a ring, these pairs are simply placed on a line. The line model follows the same basic equations as the ring model, however the stimulus input is defined differently and the recurrent connections are differently arranged.

A stimulus input is defined in terms of stimulus center *x*_0_ (taken as zero for center stimuli), length *l* and sharpness parameter *σ_RF_*. The input to an E-I pair at location *x* is given by 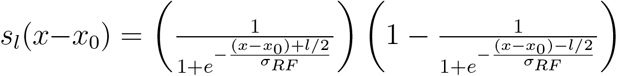. As in the ring model, this input is scaled by the overall strength of the stimulus, *c*.

In this model, there are *N* E/I units with grid spacing Δ*x*. Recurrent connections are defined with respect to distance between neurons. Excitatory projections are given by 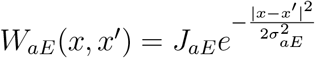 for *a* ∈ {*E*, *I*}. Inhibitory projections *W_aI_* are only to the same line position as the projecting neuron.

The parameters used in this model are: *N* = 101, 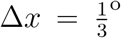, *σ_RF_* = 0.125Δ*x*, *J_EE_* = 1.0, *J_IE_* = 1.25, *W_EI_* = 1.0, *W_II_* = 0.75, 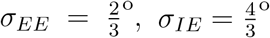, *k* = 0.01, *n* = 2.2.

Again, if multiple stimuli are present their inputs are simply added together and attention takes the same shape as a stimulus but is only directed toward E cells.

In one simulation, noise was added to the line model. This noise was similar to that added to the ring model, but with a lower baseline (5 instead of 10) and different spatiotemporal parameters: external noise was given by convolution of unit-integral Gaussian temporal filter (stdev 15 ms) and spatial filter (stdev 3Δ*x*) with Gaussian spatiotemporally white noise (mean 0, stdev 10).

#### 4.1.3. 2-D Model

The one-dimensional ring and line models vary either in preferred retino-topic location or visual feature. To create a model wherein cells have both varying retinotopic as well as feature preferences, we place E-I pairs on a two-dimensional spatial grid representing retinotopy, with an overlaid map of preferred orientation (which may be imagined to represent any circular preferred feature). This model also incorporates randomness in parameters, allowing study of diversity in responses as in Fig. 3.

Let *W_ab_*(*x,x*′) be the synaptic weight from the cell of type *b* (E or I), at position *x*′, with preferred orientation *θ*(*x*′), to the cell of type *a*, at position *x*, with preferred orientation *θ*(*x*). Nonzero connections are sparse and chosen randomly, with probability 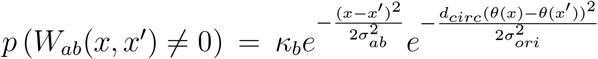. Where a nonzero connection exists, *W_ab_*(*x,x*′) is chosen randomly from a Gaussian distribution with mean *J_ab_* and standard deviation 0.25 *J_ab_*; weights of opposite sign to *J_ab_* are set to zero. For each cell, the set of recurrent synaptic weights of type *b* (E or I) it receives are then scaled so that all cells of a given type *a* (E or I) receive the same total inhibitory and the same total excitatory synaptic weight from the network, equal to *J_ab_* times the mean number of connections received under *p* (*W_ab_*(*x,x*′) ≠ 0). *τ_E_, τ_I_, n_E_, n_I_*, and *k* are also drawn from Gaussian distributions, with standard deviation 0.05 times the mean (parameter values below indicate means).

We use a grid of 75 × 75 E-I pairs. The preferred orientation of an E-I pair is given by a map randomly generated using the method of Ref. (Kaschube et al., 2010), (their supplemental materials, Eq. 20) with *n* =30 and 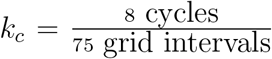. The full map is taken to be 16° × 16°; the grid interval 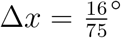. Boundaries in retinotopic space are periodic. Parameters: *κ_E_* = 0.1, *κ_I_* = 0.5, *J_EE_* = 0.10, *J_IE_* = 0.38, *J_EI_* = 0.089, *J_II_* = 0.096, *κ* = 0.012, *n_E_* = 2.0, *n_I_* = 2.2, *σ_EE_* = 8Δ*x, σ_IE_* = 12Δ*x*, *σ_EI_* = *σ_II_* = 4Δ*x, σ_ori_* = 45°, *σ_FF_* = 32°, *σ_RF_* = Δ*x*. Degrees can be converted to distance across cortex by assuming a cortical magnification factor of 0.6 mm/deg, a typical figure for 5 − 10° eccentricity in the cat (Albus, 1975) giving *σ_EE_ = σ_IE_* = 1.54mm, *σ_EI_ = σ_II_* = 0.513mm, orientation map period 1.2mm.

In this model, the relative input to the cell at 2D-position **x** with preferred orientation *θ*(**x**) from a grating of size *l* centered at position *x′* with orientation *ϕ* is 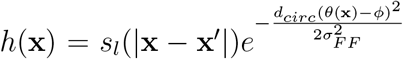; for a full-field grating, the relative input is simply 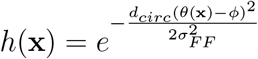.

We used different exponents, *n_I_* > *n_E_*, to increase stability despite variability (as supported by experiments: Supplemental Figure S3 of Ref. Haider et al., 2010). Variability of *τ*’s, *n*’s, *k* was limited because larger variability tended to yield instability; biologically, large variability can probably be tolerated without instability because of various forms of homeostatic compensation (Turrigiano, 2011), not modeled here.

#### 4.1.4. E-I Pair Model

In Figure 1A we study an isolated E-I pair. The inputs in this simple two-neuron model are given by:

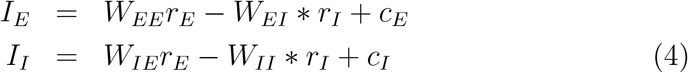

We use the following parameters: *W_EE_* = 1.00, *W_IE_* = 1.25, *W_EI_* = 0.75, *W_II_* = 0.75, *k* = 0.01, and *n* = 2.2. The inputs *c_E_* and *c_I_* are the sums of two components, an “orientation tuned” input that is equal between the two neurons and an untuned modulatory component added to either the E or I cell on a given trial. The tuned component is given by a Gaussian curve at orientation 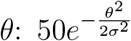, *σ* = 20°. Modulatory input: to I cells, from 0 to 10 in steps of 2.5; to E cells, from 0 to 5 in steps of 1.25.

### 4.2. Attention Experiments

Unless otherwise noted, simulations ran for 300ms and final firing rates for excitatory cells were reported. Attention was modeled as additional input of a specified strength given only to the excitatory cell in a pair. Unless otherwise stated, the shape of the attentional inputs was the same as that of the attended stimulus (as schematized in Figure 1B).

#### 4.2.1. Using the Ring Model

In Figure 2A, we used the ring model to show how attention to a non-preferred stimulus enhances suppression. The preferred stimulus was oriented at 45 degrees, with strength 40. The non-preferred was oriented at 135 degrees and the strength varied from 0 to 80. Attention was applied to either stimulus at strength 3.

In Figure 2B, a non-preferred stimulus (oriented at 135 degrees with strength 40) for the recorded cell (located at 45 degrees) was present as another stimulus (also strength 40) varied from orientation 0 to 180 degrees. Attention (strength 2) was applied to the non-preferred probe stimulus, to the varying stimulus, or not applied at all.

In Figure 5B (left), activity was recorded from a cell at 45 degrees while a preferred stimulus (45 degrees) was presented in conjunction with a non-preferred (135 degrees) stimulus. While the non-preferred stimulus remained at strength 50, the strength of the preferred one varied logarithmically from 1 to 100. Attention was directed to the non-preferred stimulus with strength 5 (or was absent). In Figure 5B (right), the contrast of both the preferred and non-preferred stimulus varied logarithmically from ≈1-20. Attention was applied either to the preferred or non-preferred stimulus with strength 1.

In Figure 7A, the cell located at 10 degrees was recorded. Each combination of a preferred stimulus (20 degrees), intermediate stimulus (60 degrees), non-preferred stimulus (80 degrees), or no stimulus was tested. All stimuli were presented with strength 20 and an additional input of 10 was given to all cells to better match the baseline firing in (Sundberg et al., 2009). Attention (of strength 1.5) was applied to either of the stimuli present or not at all.

In Figures 8A and B, the ring model with added noise was used and simulations ran for 500ms. In Figure 8A, for the first 250ms, no stimulus or attentional inputs are given (noise inputs are on throughout). At 250ms, a stimulus of strength 25 located at 90 degrees turns on, and on half of the trials so does an attentional input at the same location (strength 8). 1000 trials are run in total. To calculate spontaneous firing rates and Fano factor (FF), firing rates are averaged over 100-250ms. For stimulus-evoked activity, they are averaged over 350-500ms (these are the two epochs compared when calculating the fraction change in firing and FF in the right plot of the figure). Both E and I cells from 30-65 degrees were recorded.

In Figure 8B, for the first 250ms, no stimulus or attentional inputs are given (noise inputs are on throughout). At 250ms, two stimuli (both of strength 25, one located at 90 degrees and one at 45) turn on, and on half of the trials so does an attentional input at 90 degrees (strength 8). 1000 trials are run in total. For the figure on top, correlations are calculated in overlapping windows of 50ms for two cells (representing 25 and 30 degrees). On the bottom, correlations are calculated from firing rates averaged over 350-500ms. E cells at all locations were recorded and correlation is plotted as a function of the distance on the ring between any two pairs.

#### 4.2.2. Using the Line Model

In Figure 4A, a stimulus of strength 25 and length 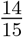 spatial degrees was either placed at the center of the receptive field of the cell at position 0, placed in its surround (at a distance of 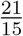 degrees), or placed at both locations simultaneously. In the last configuration, attention (strength 2) was applied either to the stimulus at the center or the surround (or not at all).

In Figure 5A (left), a stimulus of length 1 spatial degree is presented at the center of the recorded cell with contrast varying logarithmically from 1-100. Attention of strength 1 and length 25 degrees is applied at the same location. For the figure on the right, the size of the attention and stimulus are reversed. To replicate differences in baseline firing shown in (Reynolds and Heeger, 2009), an additional input of 10 is given to all cells in the simulations producing the figure on the left, and an additional input of 2 is given for those on the right.

In Figure 6A, a stimulus of strength 15 was centered on the receptive field of the recorded cell with length varying from 0 to 2.5 degrees. The size of attention (applied with strength 4) was equal to the length of the stimulus times an attention scale factor which ranged from .3 to 1.2. The preferred length is defined as the length at which the maximal firing rate is elicited.

In Figure 6B, two stimuli of contrast 15 and length 1.1 (meant to match the size of the cRF) are placed one either edge of the recorded cell’s cRF. Exact placement relative to the receptive field center is determined by drawing from a uniform distribution ranging from .2 to .49 times the cRF diameter for the stimulus on the left and .51 to .8 for the stimulus on the right (thus the left stimulus is always inside the cRF and the right stimulus always outside). A probe stimulus of the same size and contrast 30 is moved along the spatial axis in order to map the receptive field center and surround when attention (strength 3) is deployed to the left stimulus, right stimulus or not at all. The receptive field shape found by this procedure is interpolated to give a smoother measure of the location of the receptive field center, the location of the surround, and the width of the cRF (following procedures described in Anton-Erxleben et al. (2009). In total, 100 cells were stimulated.

In Figure 7B, a stimulus of length 1 degree and strength 25 is centered on the recorded neuron’s receptive field. A stimulus of the same size and strength either is or isn’t presented in the surround (1.5 degrees away). Attention (strength 1, length 1) is applied to the center or surround location in each condition.

In Figure 8C, the line model with noise added is used. Two stimuli each of length 2.75 degrees were placed at a distance of 2 degrees on either side of the center of the line model. One had a *c* of 30 and the other 65. On attention trials, attention was applied to both stimuli with a strength of 5. For each ‘recording session’ simulated, excitatory cells 39-63 (roughly 4 degrees on either side of the center cell at 51) were recorded as these cells responded to one or the other stimulus alone. Responses to each stimulus alone at *c* = 65 (50 trials each) were used to calculate a d-prime value for each cell that represents the extent to which that cell prefers one stimulus over the other. As in Ruff and Cohen (2014), the product of d-primes defined the TTS (task tuning similarity) value for a pair of cells. 100 attention trials and 100 no attention trials were run to calculate the correlation coefficients for each pair of cells in each condition based on the average firing over the final 25ms of the simulation (results are the same using 250 or 500 trials). 20 different ‘recording sessions’ were created using a different random seed for the noise with each one. In addition to the mean changes plotted in Figure 8C, we also explored the relationship between TTS and correlation by fitting separate lines to the correlation versus TTS plot in the no attention case and the attention case. If attention differently affects negative and positive TTS pairs, the slope of the attention line should be less than the no attention line. Using the same bootstrap analysis as in Ruff and Cohen (2014) we found this to be true for all 20 of our populations (not shown).

#### 4.2.3. Using the 2-D Model

In Figure 3, the two-dimensional model was used to explore the relationship between normalization and attention. We sampled 250 excitatory cells from the model. For each cell, a stimulus of preferred orientation, size 16 degrees, and strength 40 is presented to the cell. An orthogonal stimulus of the same size, position, and strength (the “null” stimulus) is then presented, and then the preferred and orthogonal stimuli are presented together. Attention (strength 8) is applied either to the preferred or null stimulus. These response values are used to calculate the normalization modulation index, defined as: NMI = [(r(Preferred) - r(Null)) - (r(Both - r(Null))]/[(r(Preferred) - r(Null)) + (r(Both - r(Null))], as well as the attention modulation index defined as: AMI = (r(Attend Preferred) - r(Attend Null))/(r(Attend Preferred) + r(Attend Null)) for each cell.

In Figure 4B, we sample 100 cells from the model to test the interaction between surround suppression and attention. For each cell, a stimulus of strength of 50 of preferred orientation and size 10 degrees is shown. A stimulus with the same orientation and strength is placed in the surround at a distance of 10 degrees, and the response is recorded. The surround at 10 degrees is, technically, a circumference of possible positions around the center. To decide where to place the surround stimulus, the surrounding neuron at a distance of 10 with a preferred orientation closest to that of the center neuron is chosen. Attention (modulation strength = 5) is then directed either to the center or surround stimulus.

### 4.3. The SSN-CNN Model and Experiments

The SSN-CNN is an adaptation of a traditional convolutional neural network. The inputs to the network are grayscale images of handwritten digits (28-by-28 pixels). The first convolutional layer applies 180 separate 3×3 filters, all of which are constrained during training to contain only non-negative values. The application of these filters results in 180 feature maps, each with a spatial dimension of 28× 28. A 3× 3 max-pooling layer with stride 2× 2 reduces the feature map size down to 14× 14. The output of the pooling layer determines the input to the ring SSNs that exist at the next layer. Specifically, at each of the locations on the 14×14 spatial map, there is a ring SSN with 180 E/I pairs. The activity of the units in the 180 feature maps provide the *c* values (that is, the strength) for inputs centered at that location on the ring. We arbitrarily number the feature maps from 1 to 180 and let *ϕ* be the number of a particular feature map. Then at spatial position *x,y*, the feedforward input to each cell in the E-I pair located at position *θ* in the ring model is given by ∑_*ϕ*_ *c_xy_* (*ϕ*) *h*(*θ,ϕ*), with *c_xy_*(*ϕ*) the activity of the unit in the *ϕ* feature map in the pooling layer at location *x,y*, and *h*(*θ,ϕ*) the function defined in section 4.1.1. While there is no concept of a ring in the topology of the feature maps prior to learning, we still map the 180 feature maps onto the 180 locations in the ring. Because feature maps assigned to more nearby locations in the ring will more strongly influence one another’s output on the ring, the feature maps should ultimately develop structure reflecting the ring topology (Lindsay and Miller, 2018).

This architecture is then repeated to create a two-layer convolutional network. The output of the second SSN layer serves as input to a fully-connected layer with 1024 units, which then projects to the final 10-unit layer (one for each digit). For training, the network was unrolled for 46 timesteps (with *dt* = 2ms for the SSN layers) and trained on the MNIST dataset using backpropagation through time to minimize a cross entropy loss function (batch size 128). Only the final timestep was used for calculating the loss function and classification accuracy. The recurrent weights for each ring SSN at both layers were set as described above for the standard ring network. These weights were not allowed to change during training.

Repeating the procedure of (Lindsay and Miller, 2018), once the network was trained on the standard classification task, the final 10-unit layer was replaced with a series of binary classifiers, one for each digit. The weights from the 1024-unit second-to-last layer to the 2-unit final layer were trained to perform binary classification on a balanced training set wherein half of the images were of the given digit and half without.

We then generate more challenging images on which to test the benefits of attention. These images consist of two regular MNIST images added together. The test set for each binary classifier contains 768 images, half of which contain (as one of the two digits) the digit the classifier was trained to detect and the other half do not. Performance accuracy is given as the overall percent correct of the binary classifier on this test set.

To know how to apply attention, we first present 45 standard MNIST images of each digit to the network and record the activity of neurons in the SSN. From this we calculate “tuning values” that indicate the extent to which each feature map prefers each digit. As in (Lindsay and Miller, 2018), tuning values are defined as a z-scored measure of the feature map’s mean response to each digit. Specifically, for feature map *θ* in the *l^th^* layer, we define *r^l^*(*θ, n*) as the activity in response to image n, averaged over all units in the feature map (i.e., over the spatial dimensions). Averaging these values over all images in the training sets (*N_d_* = 45 images per digits, 10 digits. N=450) gives the mean activity of the feature map 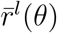:

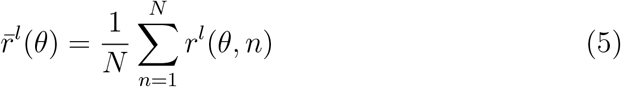

Tuning values are defined for each feature map and digit, *d* as:

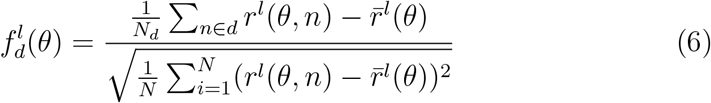

When attention is applied to a particular digit, excitatory neurons that prefer that digit are given additional input. Specifically, the cells in feature maps whose tuning value for the attended digit are above the median tuning value for that digit are given attentional inputs. The attentional input to each feature map is proportional to how much above the median its tuning value is:

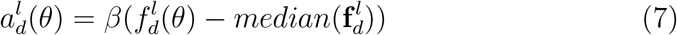

Note, in this model the attentional input to the excitatory cell is fully specified by the above equation (that is, this value is not multiplied by the shape of the feedforward input).

We define digit preference on the feature map level (rather than for individual neurons) because feature attention is known to be a spatially-global phenomenon (that is, attention applied to a particular feature modulates neurons at all spatial locations, (Saenz et al., 2002)).

The accuracy on the same test set of overlaid images is again calculated for each digit, now in the presence of attention directed to the digit being detected. An additional parameter representing the overall strength of attention (*β*) is varied (.02, .04, or .06) and for each digit the best performing strength is used.

This attention was applied at each SSN layer individually as well as at both together. Here, the results of applying attention at the second SSN layer are reported as this elicited the best performance (a finding that is in line with those reported in (Lindsay, 2015; Lindsay and Miller, 2018), wherein attention at later layers better enhanced performance).

## 5. Acknowledgements

We thank Daniel Bear, Aran Nayebi, and other members of Daniel Yamins’s lab for help with the code used to train the SSN-CNN. This work was supported by funding from the Gatsby Foundation, Google, National Science Foundation (NeuroNex DBI-1707398 and IIS-1704938), and the Simons Collaboration on the Global Brain (543017).

## Appendix A. Supplementary Figures

**Figure A.11:**
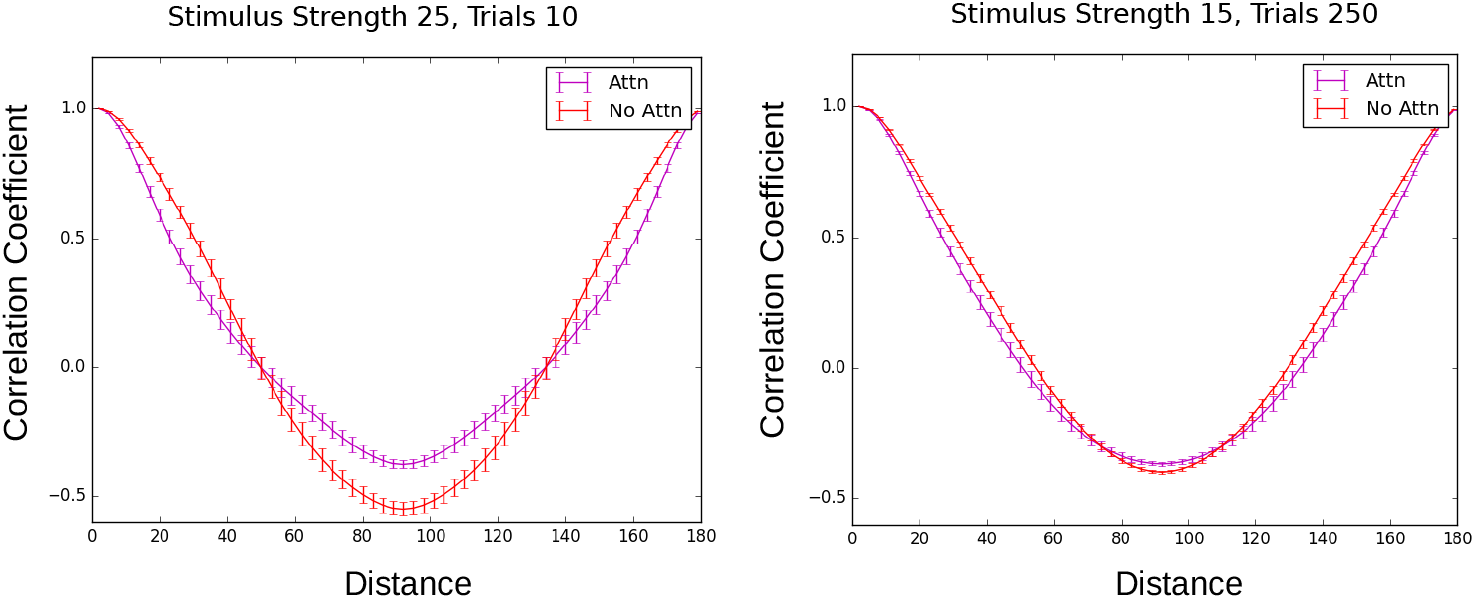
Attention can increase correlations. Example runs of the model used to make Figure 8B that result in attention increasing correlations for distant pairs. The strength of the stimulus and number of trials used for each condition is given at the top for each (in Figure 8B, strength was 25 and 500 trials were used). Errorbars are SEM.

**Figure A.12:**
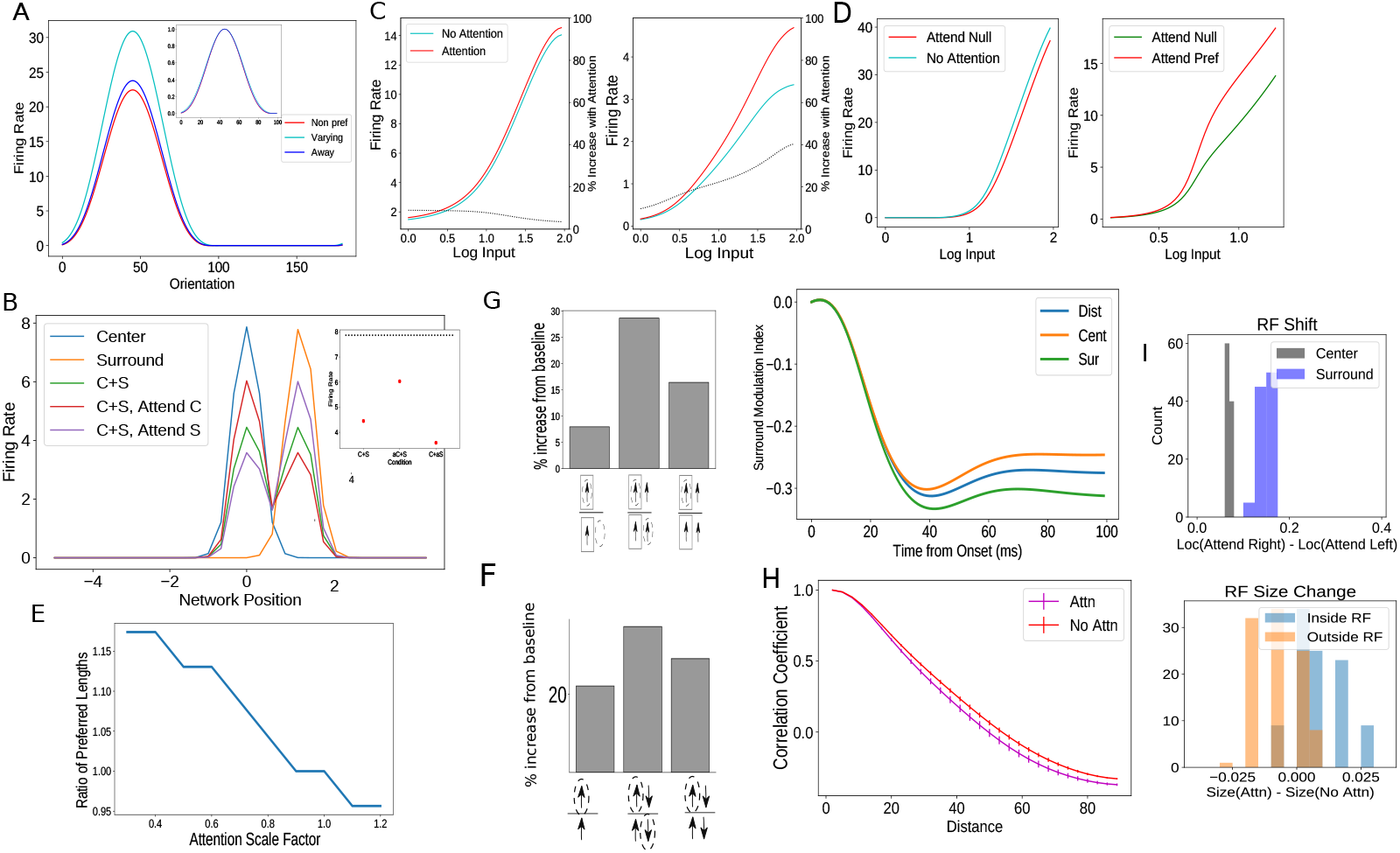
Findings that qualitatively replicated with attention modeled as inhibitory input to inhibitory cells. A. Replication of Figure 2B. B. Replication of Figure 4A. C. Replication of Figure 5A. D. Replication of Figure 5B. E. Replication of Figure 6A. F. Replication of Figure 7A. G. Replication of Figure 7B. H. Replication of Figure 8B. I. Replication of Figure 6B.

**Figure A.13:**
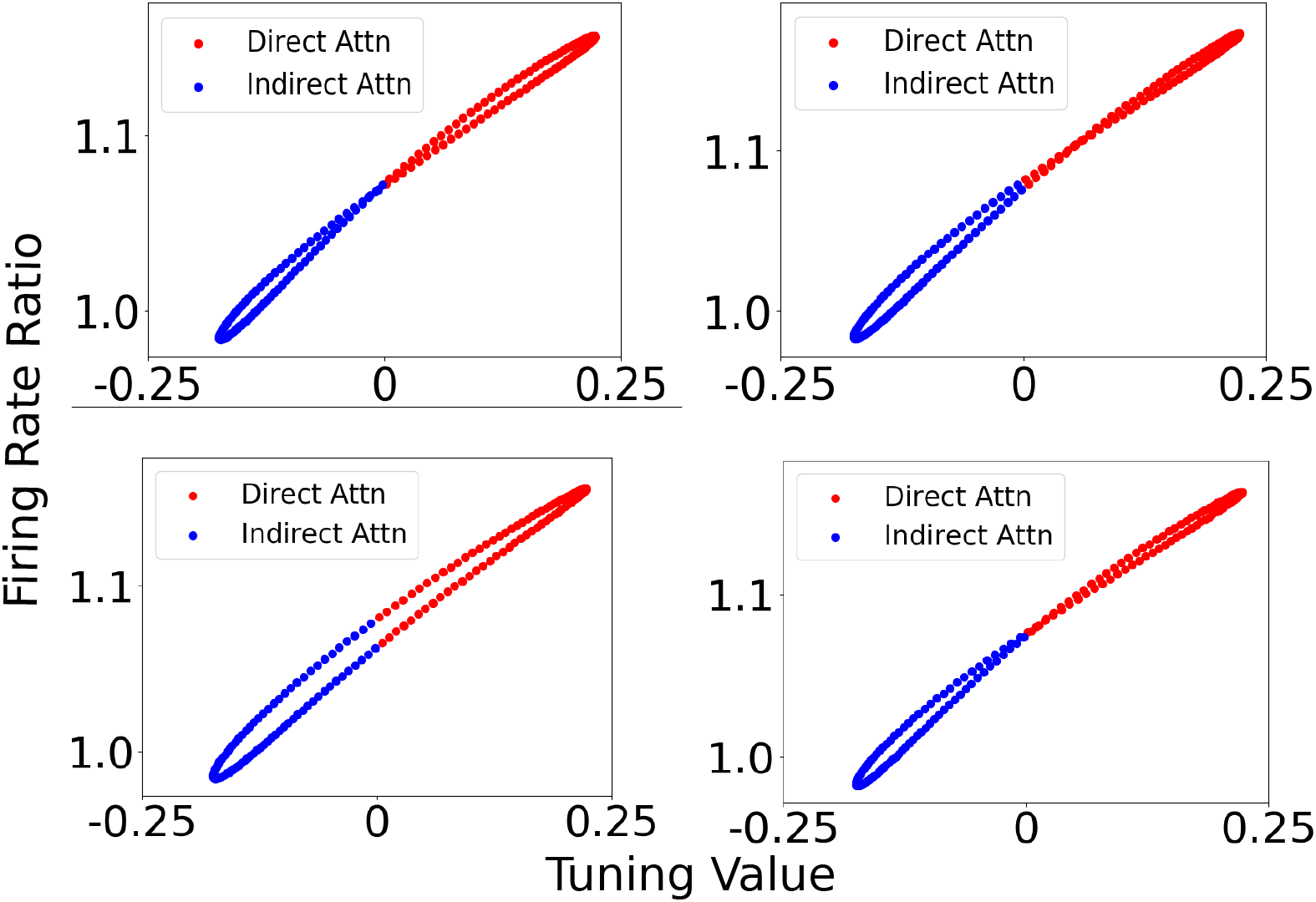
Impact of feature attention at different spatial locations in layer 2 of the SSN-CNN. Ratio of attended to non-attended firing rates for cells in a ring network as a function of tuning value as in Figure 10E, but for different nearby spatial locations.

